# Length-dependent hypercontractility decoupled from cellular adaptations is the initial phenotype in MYBPC3-haploinsufficient iPSC-derived engineered cardiac tissues

**DOI:** 10.64898/2026.05.13.724801

**Authors:** Tomi Tuomainen, Maike Mona Shirinov, Mina Rabiee, Eloi Schmauch, Jussi T. Koivumäki, Kyriakitsa Galani, Manolis Kellis, Johanna Kuusisto, Robert S. Leigh, Nikolay Naumenko, Marko Vendelin, Suvi Linna-Kuosmanen, Pasi Tavi

## Abstract

Hypertrophic cardiomyopathy (HCM) is often characterized by a complex landscape of secondary compensatory changes, making it difficult to distinguish primary events triggered by the underlying genetic variants. Here, using patient-derived iPSC-engineered heart tissues cultured in conditions minimizing external stimuli, we identify the initial phenotype of the HCM inducing variant MYBPC3-Q1061X. We demonstrate that initially MYBPC3 haploinsufficiency leads to robust, muscle length-dependent hypercontractility with emerging diastolic dysfunction at higher contraction rates. This phenotype is entirely decoupled from early transcriptional and metabolic stress responses, as evidenced by physiological phenotype and metabolite and cell type-specific transcriptional profiles. Mathematical modeling reveals that the hypercontractile phenotype can be translated to adult cardiac muscle with a small change in myosin availability, amplified by inherent cooperativity within the sarcomere that is instrumental in enhancing the force and the speed of the contraction cycle. Despite the apparent calcium sensitization, these sarcomeric changes do not affect excitation-contraction coupling or calcium buffering in this proximal stage. These results suggest that the primary biophysical defect in MYBPC3-haploinsufficiency manifests as a latent mechanical hypersensitivity that precedes secondary cellular remodeling and functional instability. This initial phenotype provides a window for preventive therapeutic intervention before the onset of permanent cellular remodeling in the heart.

## Introduction

Hypertrophic cardiomyopathy (HCM) is the most common inherited cardiomyopathy in adults, affecting approximately 1 in 500 individuals (Maron et al., 2012). More than 1,500 pathogenic variants have been identified, primarily in sarcomeric and Z-disc genes (Maron et al., 2014).

Mutations in the myosin binding protein C3 (*MYBPC3*) gene are the most frequent accounting for over 40% of all familial HCM cases (Chiswell et al., 2023), highlighting *MYBPC3* as a major genetic driver of HCM. Over two-thirds of pathogenic *MYBPC3* variants are truncating (frameshift or nonsense), typically leading to absent or unstable protein products (Barefield et al., 2014; Helms et al., 2014), supporting the widely accepted notion that haploinsufficiency is the primary disease mechanism (Barefield et al., 2015; Glazier et al., 2019; Helms et al., 2014; Marston et al., 2009; van Dijk et al., 2009). However, regulation of cardiac myosin binding protein C (cMyBP-C) is complex, as reduced protein production can be offset by decreased degradation, yielding near-normal cMyBP-C levels *in vitro* (Helms et al., 2020), and genetic predisposition for haploinsufficiency may manifest only under stress (Barefield et al., 2015).

cMyBP-C is a key structural and regulatory component of the sarcomere, that links thin and thick filaments (Luther et al., 2011), thereby establishing interfilamentous regulation (Herron et al., 2006). The *MYBPC3* gene spans >21 kb and 34 exons (Carrier et al., 1997) encoding a protein with binding sites for both thin and thick filaments in its N- and C-terminal domains (Carrier et al., 2015; Harris, 2021). Functionally, cMyBP-C restricts the actin–myosin interactions, thereby regulating force generation (Gruen and Gautel, 1999; Hartzell, 1985; Kensler et al., 2011; Sarkar et al., 2020). Loss of cMyBP-C accelerates cross-bridge kinetics, increases energy demand, and alters sarcomere mechanics (Desjardins et al., 2012; Korte et al., 2003; Palmer et al., 2004). Cardiac MyBP-C also stabilizes a subpopulation of myosin heads in the “super-relaxed state” (SRX), a low ATP-consuming reserve that can be mobilized by stretch, phosphorylation, or truncating *MYBPC3* mutations (Ma et al., 2021; McNamara et al., 2017; McNamara et al., 2016; McNamara et al., 2019). These regulatory effects are fine-tuned by post-translational modifications, particularly phosphorylation by kinases downstream of adrenergic signaling, which coordinate force generation, relaxation, and length-dependent activation (Gupta and Robbins, 2014; Hanft et al., 2021; Kensler et al., 2017; Moss et al., 2015).

Although cMyBP-C is dispensable during embryonic heart development (Harris et al., 2002), it is essential for maintaining sarcomere integrity and function in adulthood (Carrier et al., 2015). Mice lacking cMyBP-C (*Mybpc3-/-*) are viable at birth but develop hypertrophy at a young age, which is associated with increased myofilament Ca²⁺ sensitivity, contractile dysfunction, and mitochondrial abnormalities (Harris et al., 2002). Heterozygous knock-in mice carrying the HCM-causing *MYBPC3*-G5256A variant (Richard et al., 2003) show diastolic dysfunction and myofilament Ca²⁺ sensitization prior to hypertrophy, suggesting that haploinsufficiency initiates subtle functional changes that precede pathology (Fraysse et al., 2012). In patients, hypercontractility due to sarcomeric mutations often manifests as an elevated left ventricular (LV) ejection fraction before hypertrophy develops (Captur et al., 2014; Ho et al., 2002). However, patient-derived myocardial preparations show heterogeneous phenotypes, including impaired relaxation, altered Ca²⁺ sensitivity, reduced maximal force, and energetically costly hypercontractility (Jacques et al., 2008; Pioner et al., 2023; van Dijk et al., 2012). In human iPSC-CMs, HCM-inducing MYBPC3 mutations induce a variable spectrum of features typically associated with advanced disease, ranging from sarcomeric disarray to dysregulated Ca²⁺ signaling and electrophysiological abnormalities (Li et al., 2022; Mori et al., 2024; Ojala et al., 2016; Prondzynski et al., 2017). In more mature and functionally competent iPSC-derived engineered heart muscles (Eder et al., 2016) cMyBP-C haploinsufficiency initially leads to a hypercontractile phenotype, followed by the development of contractile dysfunction and calcium signaling abnormalities (De Lange et al., 2023).

The central unresolved questions regarding most HCM mutations pertain to understanding the function of the proteins involved, which is a prerequisite for understanding how their perturbation leads to cardiac remodeling. While the sequence of events linking *MYBPC3* mutations to HCM remains not fully understood, it is apparent that pathogenic variants trigger early molecular and functional changes that are often exacerbated by additional somatic stressors (Wolf et al., 2005) to gradually evolve into hypertrophy. To characterize the biophysical basis of the initial phenotype, before secondary HCM remodeling, we focused on the most common HCM mutation in Finland: MYBPC3-Q1061X, a founder variant introducing a premature stop codon in exon 29 that truncates the C-terminus (Jaaskelainen et al., 2002). The resulting cMyBP-C haploinsufficiency causes HCM with a relatively late onset and ∼78% penetrance (Jaaskelainen et al., 2013; Jaaskelainen et al., 2004). Here, we define physiological changes in engineered heart tissues (EHTs) derived from patient iPSC lines carrying Q1061X and isogenic controls. Assessment of the functional changes is combined with single-cell transcriptomics and mass spectrometry-based metabolomics in search of common HCM remodeling signs. Computational modeling of the adult sarcomere recapitulates the iPSC-EHT phenotypes, identifies molecular mechanisms underlying the earliest consequences of MYBPC3 haploinsufficiency, and provides insight into the primary interactions initiating human HCM.

## Results

### MYBPC3-Q1061X causes hypercontractility but not hypertrophy in patient specific iPSC-derived EHTs

iPSC lines originating from two patients carrying MYBPC3-Q1061X-mutations (Fig. 1A) and their corresponding gene-edited isogenic control lines produced comparable amounts of TNNT2-positive cardiomyocytes after standard cardiac differentiation (Fig. 1B). After casting and culturing under minimal loading conditions by keeping the muscles near slack length these lines produced comparably sized engineered heart tissues (EHTs) (Fig. 1C,D) with equally sized cardiomyocytes (Fig. 1E). MYBPC3-Q1061X mutation is predicted to induce truncation of the protein leading to MyBP-C haploinsufficiency (Jorgenrud et al., 2015), which was confirmed in tissues carrying the mutation (Fig. 1F). Electrically evoked (1 Hz) isometric force was significantly higher in EHTs harboring MyPB-C mutation than in their corresponding controls (Fig. 1G) indicating a hypercontractile phenotype. Single-nuclei RNA sequencing (snRNA-seq) identified major cardiac cell types from EHTs, including cardiomyocytes, fibroblasts, endothelial cells, mesothelial cells, and a population with transcriptional profiles characteristic of developing cells (Fig. 1H). MYBPC3 expression was specific to cardiomyocytes (Fig. 1I), consistent with other cell type-specific markers that delineate distinct identities (Fig. 1J). The integration of our dataset with previously published adult human left ventricle snRNA-seq data (Chaffin et al., 2022) showed that while EHT cell clusters largely overlap with their adult counterparts, they do not capture the full spectrum of adult cell transcriptional variability (Fig. 1H). These results demonstrate that in iPSC-derived EHTs containing multiple cardiac cell types, MyBP-C haploinsufficiency in cardiomyocytes leads to tissue hypercontractility without hypertrophy at the tissue or cellular level.

**Figure 1.**
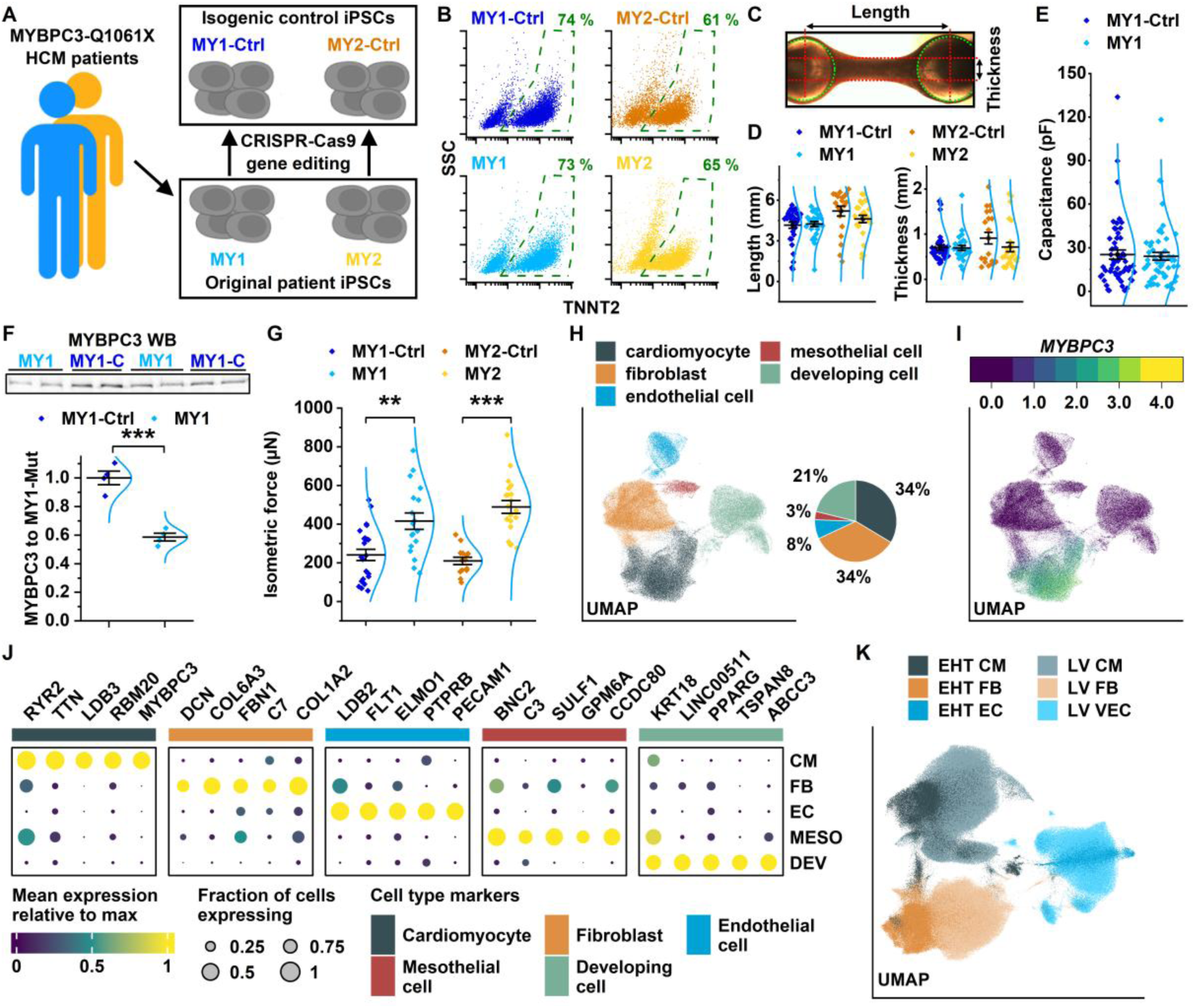
Characterization of *in vitro* cardiac MYBPC3-Q1061X engineered heart tissue model. **A)** Schematic representation of the production of iPSC lines used in the present study. **B)** Representative flow cytometric analysis showing the fraction of cardiac troponin T (TNNT2) positive cells in 2D iPSC-CM cultures prior to the production of engineered heart tissues (EHT). Dashed green line highlights the TNNT2 positive cell fraction in each scatter plot. **C)** Representative image of the EHT during force recording. Dashed green line shows the position of PDMS rack post ends and dashed red lines illustrate how length and thickness of the EHT were determined from the image. **D)** Length (*left*) and thickness (*right*) of the EHTs at the time of force recording. n(MY1-Ctrl)=32, n(MY1)=28, n(MY2-Ctrl)=20, n(MY2)=25. **E)** Cell membrane capacitance assessed from individual cardiomyocytes after dissociation and plating of cells cultured in EHTs. n(MY1-Ctrl)=54, n(MY1)=57. **F)** Western blot analysis of MYBPC3 protein in EHTs. n=4. **G)** Twitch force produced by EHTs during isometric force recording. n(MY1-Ctrl)=23, n(MY1)=18, n(MY2-Ctrl)=14, n(MY2)=20. **H)** Uniform manifold approximation and projection (UMAP) plot of single-nuclei RNA sequencing (snRNA-seq) analysis of EHTs showing the annotation of the five main cell types and their proportions in the whole snRNA dataset. **I)** UMAP plot showing expression level of *MYBPC3* in EHT nuclei. **J)** Relative expression levels and fractions of cells expressing the selected cell-type markers in different cell types of EHTs. **K)** UMAP showing the overlap of cell types shared by EHTs and human left ventricular snRNA-seq data after dataset integration. In F) Student’s *t*-test *P* value: ***, *P* < 0.001. In G) ANOVA with Bonferroni *post hoc* test: **, *P* < 0.01; ***, *P* < 0.001

### MYBPC3-Q1061X mutation does not cause HCM-associated transcriptional or metabolic remodeling in EHTs

The central hallmarks of HCM include transcriptional (Wehrens et al., 2022) and metabolic remodeling (Nollet et al., 2024), which drive disease progression. To determine whether MYBPC3-Q1061X induces similar changes in EHTs, cell type-specific transcriptional profiles were analyzed using snRNA-seq and metabolic alterations by using untargeted LC-MS metabolomics. Cardiomyocytes from MYBPC3-Q1061X EHTs exhibited 221 altered transcripts (Fig. 2A), mostly associated with pathways unrelated to HCM such as cardiogenesis and neuron-related functions (Fig. 2B). Some of these transcripts were minimally expressed in adult cardiomyocytes or were changed to opposite direction in response to disease variant compared to HCM-induced changes in adult cardiomyocytes (Fig. 2C). Other EHT cell types showed no clear cell-type specific transcriptional changes in response to MYBPC3-Q1061X variant (Fig. S1). Unsupervised clustering of genes related to excitation-contraction coupling (Fig. 2D) or energy metabolism (Fig. 2E, Fig. S2A) revealed distinct transcriptional profiles between EHT and adult heart cardiomyocytes. Importantly, MYBPC3-Q1061X EHTs showed no signs of HCM-associated remodeling (Chaffin et al., 2022). Although energy metabolism pathways were profoundly remodeled in HCM hearts, transcriptional changes among genes related to glycolysis and β-oxidation were negligible in MYBPC3-Q1061X EHTs compared to those in adult HCM hearts (Fig. 2E, Fig. S2A). Consistent with this, while untargeted LC-MS analysis comparing MYBPC3-Q1061X EHTs with isogenic controls identified numerous metabolites (Fig. 2F) there were no patterns resembling those observed in human HCM myocardium (Previs et al., 2022) (Fig. 2G). Furthermore, both respiratory and glycolytic capacities of isolated cardiomyocytes from MYBPC3-Q1061X EHTs were unchanged compared to those of the controls (Fig. 2H,I, Fig. S2B,C). Together, these findings indicate that the MYBPC3-Q1061X mutation does not induce HCM-associated transcriptional or metabolic remodeling in iPSC-derived EHTs, which further suggests that additional stimuli are prerequisite for remodeling.

**Figure 2.**
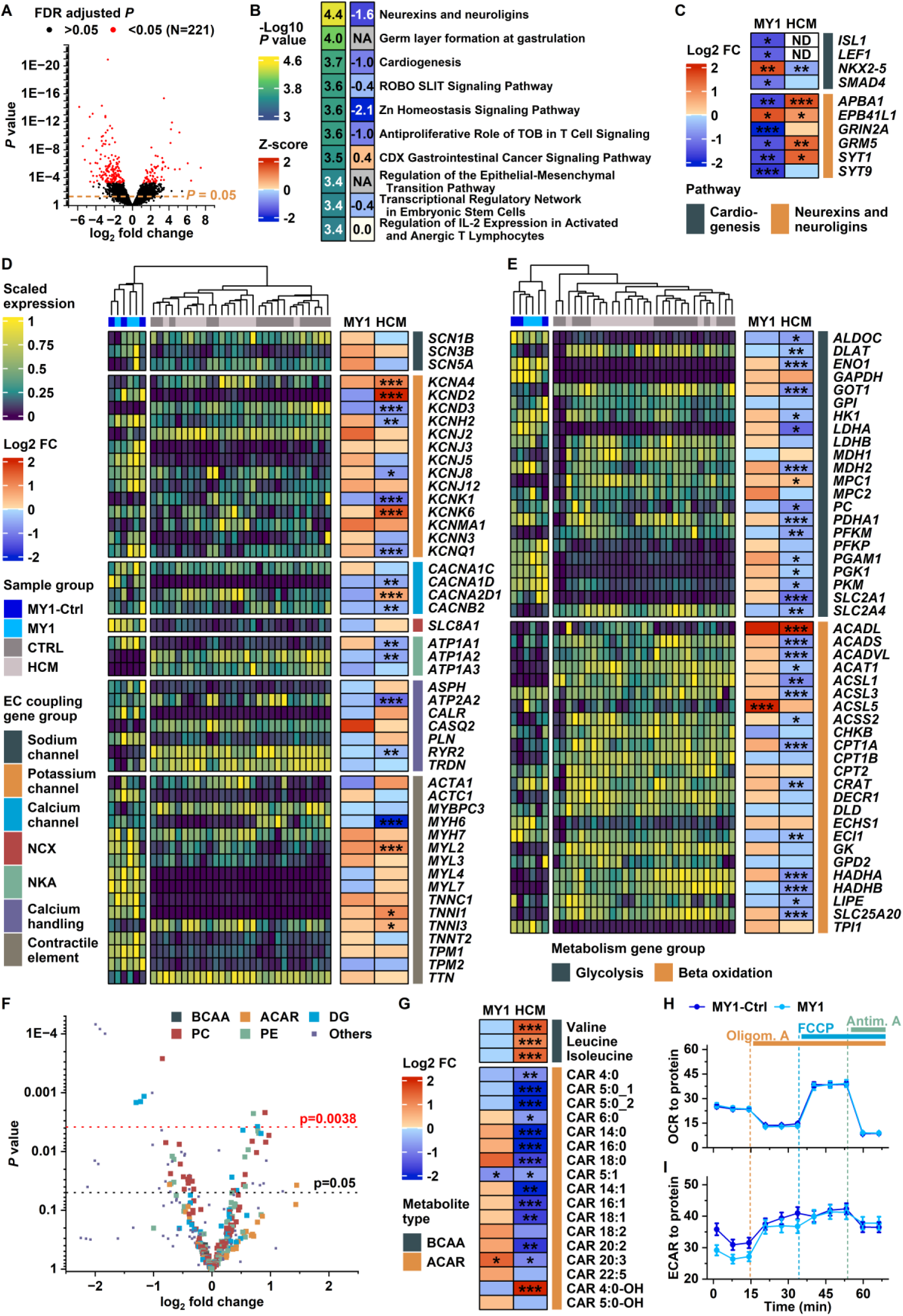
Transcriptional and metabolic changes typical for hypertrophic cardiomyopathy are not induced MYBPC3-Q1061X EHT model. **A)** Scatter plot of adjusted *P* value and logarithmized fold change for each gene acquired from differential expression analysis of MY1 and MY1-Ctrl EHT cardiomyocytes from snRNA-seq data. n=3 (tissues in experimental group). **B)** Heatmap showing negative of logarithmized *P* value and Z-score for top 10 enriched pathways (according to enrichment *P* value) in DE genes between MY1 and MY1-Ctrl cardiomyocytes (FDR adjusted *P* value below 0.05 in panel A). **C)** Comparison of transcriptional changes (logarithmized fold change) between cardiomyocytes of EHTs (MY1) and left ventricles of hypertrophic cardiomyopathy patients (HCM) (Chaffin et al., 2022) relative to MY1-Ctrl and healthy individuals, respectively, in the genes enriched in selected pathways from panel B. **D)-E)** Comparison of excitation-contraction coupling (D) and metabolism (E) gene expression in cardiomyocytes of EHT model and HCM left ventricular tissue. Gene expression levels in individual samples are shown on the left, and differential expression on the right side of the heatmap. Hierarchical clustering was applied to the columns of the expression-level heatmap. **F)** Scatter plot of *P* value plotted against logarithmized fold change from comparison of metabolite abundance between MY1-Ctrl and MY1 EHTs. The adjusted α level for differential metabolite abundance is highlighted in red. n=10. **G)** Comparison of changes in branched-chain amino acid (BCAA) and acylcarnitine (ACAR) abundances in MY1 EHTs and heart tissue of HCM patients (fold changes and *P* values for HCM acquired from previously published data (Previs et al., 2022)). **H)-I)** Average oxygen consumption rates (H) and extracellular acidification rates (I) from mitochondrial stress test. n(MY1-Ctrl)=22, n(MY1)=24. In C), D) and E) FDR adjusted *P* value from *DESeq2*: *, *P* < 0.05; **, *P* < 0.01; ***, *P* < 0.001. In G) for MY1 Student’s *t*-test *P* value: *, *P* < 0.05, and for HCM previously assessed *P* value: *, *P* < 0.05; **, *P* < 0.01; ***, *P* < 0.001.

### MYBPC3-Q1061X-induced hypercontractile phenotype modulates muscle length-dependent activation

Phenotypes in hypertrophic cardiomyopathy (HCM) and MyBP-C ablation are commonly associated with alterations in cardiomyocyte electrophysiological properties and calcium signaling (Birket et al., 2015; De Lange et al., 2023; Pioner et al., 2023). However, confocal Ca^2+^ imaging and whole-cell voltage clamp experiments showed that MYBPC3-Q1061X-carrying myocytes isolated from EHTs exhibited no changes in action potential waveform, resting membrane potential, calcium transients, sarcoplasmic reticulum Ca^2+^ load, or L-type calcium current properties (Fig. S3) which could explain or significantly contribute to the observed hypercontractility. This suggests that hypercontractility induced by the MYBPC3-Q1061X mutation likely arises from altered regulation of contractile elements due to MyBP-C loss. While all EHTs studied exhibited physiological length-dependent activation (LDA) i.e. Frank-Starling responses, EHTs derived from MYBPC3-Q1061X lines generated higher isometric force at longer muscle lengths compared to controls (Fig. 3A,B) with significant interaction between muscle length and Q1061X-mutation in isogenic cell line pairs (Table S1). In parallel with augmented force, the mutation also increased rates of force development and relaxation, with greater effects at longer muscle lengths (Fig. 3C-E, Table S1). Although the mutation greatly enhanced force development, it did not affect passive force or viscoelastic properties of EHTs (Fig. S4).

**Figure 3.**
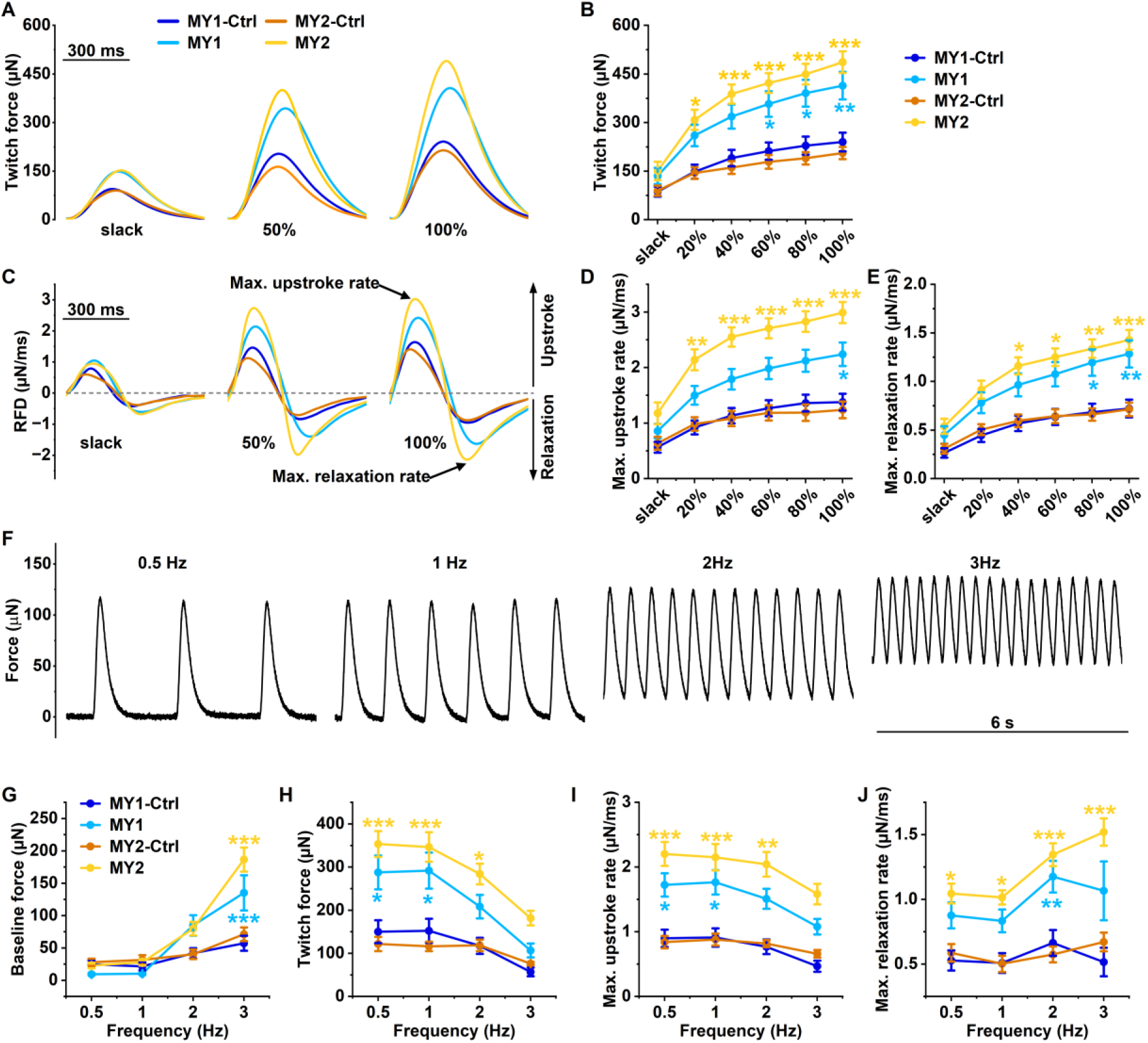
Contractile phenotype of the MYBPC3-Q1061X EHT model. **A)** Representative traces of isometric contraction twitch force at different muscle lengths, relative to length at maximal (100%) developed force for each tissue. **B)** Twitch force assessed from the isometric contraction at different muscle lengths. n(MY1-Ctrl)=32, n(MY1)=28, n(MY2-Ctrl)=20, n(MY2)=25. **C)** Rate of force development (RFD) during isometric twitch at different muscle lengths acquired by derivatization of isometric contraction traces in panel A. **D)-E)** Isometric contraction upstroke (D) and relaxation (E) rates at different muscle lengths determined from RFD plots as highlighted in panel C. **F)** Representative isometric force trace from pacing frequency protocol. **G)-J)** Baseline force (G), twitch force (H), upstroke (I) and relaxation (J) rates assessed from the pacing frequency protocol force recordings. In (B), (D) and (E), n(MY1-Ctrl)=32, n(MY1)=28, n(MY2-Ctrl)=20, n(MY2)=25. In (G-J), n(MY1-Ctrl)=7-19, n(MY1)=5-18, n(MY2-Ctrl)=7-11, and n(MY2)=12-17. *P* value of linear mixed effect model contrast between Q1061X carrying and corresponding isogenic control EHTs: *, *P* < 0.05; **, *P* < 0.01; ***, *P* < 0.001

Next, the impact of hypercontractility on other physiological features was assessed by examining the force-frequency relationship (FFR). EHT lengths were adjusted to produce 50% of maximal force before stimulation at increasing frequencies from 0.5 to 3 Hz (Fig. 3F). With increasing frequency, MYBPC3-Q1061X EHTs showed a significant increase in baseline force, reduced twitch force, and moderate slowing of contraction upstroke velocity compared to control EHTs (Fig. 3G-I, Table S2). Despite a moderate increase in relaxation velocity (Fig. 3J, Table S2), hypercontractility resulted in incomplete relaxation manifested as elevated basal tension, and reduced force production at high frequencies, consistent with diastolic dysfunction reported in MyBP-C haploinsufficient mouse models (Fraysse et al., 2012). These findings demonstrate that in iPSC-derived EHTs MyBP-C haploinsufficiency induces muscle length-dependent hypercontractility which modulates frequency response and may compromise diastolic function.

### Increased myosin availability and sarcomeric cooperativity upon loss of MyBP-C recapitulates hypercontractility *in silico*

To elucidate the fundamental drivers of hypercontractility, their collective impact on energetic costs within the pre-pathological hypercontractile phenotype, and how our results might translate to adult tissue, we employed advanced mathematical modeling of the adult sarcomere and investigated how MyBP-C haploinsufficiency modulates cooperativity, myosin ON/OFF transitions, and calcium sensitivity. The modelling platform developed here included the primary effects of MyBP-C on sarcomere function, stabilization of the super-relaxed (SRX) state and physical interfilamentous inhibition, both of which lead to a net decrease in active cross-bridge formation (Fig. 4A). Importantly, any change in the number of active cross-bridges, including that induced by altered MyBP-C binding (Heling et al., 2020; Kampourakis et al., 2014), will be amplified in speed and amount for force development by cooperativity among contractile processes (Solaro, 2010). To account for that we used a mathematical model of the sarcomere with cooperative interaction between cross-bridges and regulatory components in thermodynamic consistency between simulated reactions and developed force (Kalda and Vendelin, 2020).

**Figure 4.**
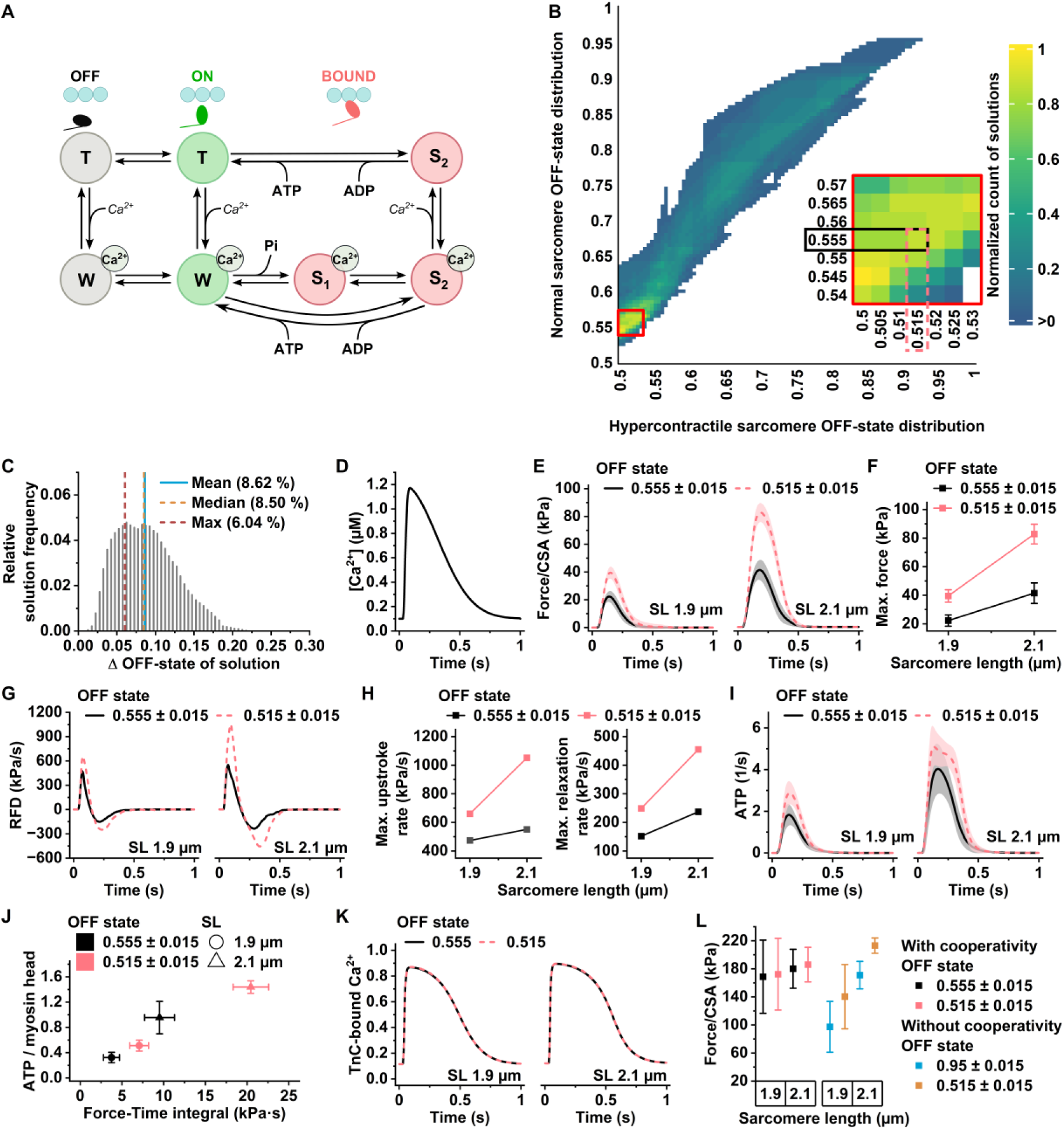
Mathematical modeling recapitulates the hypercontractile phenotype. **A)** Schematic of the contraction model. The model comprises distinct actomyosin biochemical states. In the T state, the myosin binding site on actin is blocked by tropomyosin, whereas in the W state, calcium binding to troponin C (TnC) exposes the site. These thin filament states can occur regardless of whether myosin is in the OFF or ON state, with the rates of calcium-binding transitions being identical in both myosin states. Strongly bound states (S1Ca, S2Ca, and S2) represent conformations in which myosin is firmly attached to actin and capable of generating force, differing in their free energy profiles. All transitions are reversible, with the ratio of forward to backward rates determined by the free energy difference between the corresponding states; these rates and energy parameters were optimized within the model. Transitions between the OFF and ON states are governed by an equilibrium constant and are assumed to occur on a much faster timescale than all other transitions. **B)** Heatmap of myosin OFF-state distributions in normal and hypercontractile sarcomeres. Color indicates the normalized number of model solutions for each distribution combination that reproduces the force changes observed in EHT experiments. Inset highlights the area of proportions yielding the highest density of solutions; individual combinations selected as a representative example in subsequent panels are indicated. **C)** Distribution of myosin OFF-state differences across solutions. It should be noted that most of the solutions lie in the lower range. **D**) Synthetic Ca^2+^-transient used as input to the model. **E)** Isometric twitch forces at two sarcomere lengths. **F)** Maximum isometric forces at two sarcomere lengths, obtained from panel E. **G)** Rate of force development (RFD) during isometric twitch at different sarcomere lengths acquired by derivatization of isometric contraction traces in panel E. **H)** Maximum isometric contraction upstroke and relaxation rates at different sarcomere lengths determined from RFD plots in panel G. **I)** ATP consumption rate per myosin at two sarcomere lengths. **J)** Total ATP consumed per myosin over a single cardiac cycle and corresponding force–time integral at two sarcomere lengths. **K)** Proportion of regulatory units with Ca²⁺ bound to TnC. **L)** Steady-state forces at two sarcomere lengths for conditions with and without cooperativity. Results are presented as mean ± SD.

To determine whether MYBPC3-Q1061X hypercontractility can be mechanistically reproduced by a shift in myosin head availability, simulations were performed in which only the equilibrium between ON (recruited for force generation) and OFF (super-relaxed) myosin states was varied, while all other model parameters were held constant. For simplicity, this equilibrium was treated as sarcomere length independent. Model parameters were first established by fitting adult trabeculae tension development and ATPase activity data from published data (see Methods), then adjusted for the iPSC-EHT context by shifting the baseline ON/OFF distribution toward a larger OFF-state contribution relative to adult tissue, consistent with lower myosin availability in less mature iPSC-derived cardiomyocytes (Walklate et al., 2022). The MYBPC3-Q1061X mutation was then modeled as a shift of this equilibrium back toward the ON state, and parameter combinations were accepted when the simulated mutant-to-control force ratio matched which was measured experimentally. Because many parameter sets fit the control data equally well, simulations were conducted across a large ensemble of parameter combinations, enabling results to be interpreted as changes in parameter distributions rather than single-point estimates.

The ensemble approach revealed a broad but clustered set of ON/OFF equilibrium combinations capable of reproducing the observed changes in contractility (Fig. 4B). Across accepted solutions, the mutation shifted the ON/OFF equilibrium by ∼8% on average, with the highest solution density near 4% (Fig. 4B inset, Fig. 4C). These solutions were obtained under the constraint that the cytosolic Ca²⁺ transient remained unchanged (Fig. 4D), and representative examples confirmed that the model accurately reproduces both the magnitude and rate of force development observed in MYBPC3-Q1061X EHTs (Fig. 4E–F). Consistent with the increased force-time integral, the mutation was predicted to elevate ATPase rate and net ATP consumption per twitch (Fig. 4I,J). Notably, troponin C-bound calcium remained unchanged across solutions (Fig. 4K), indicating that the apparent increase in myofilament Ca²⁺ sensitivity arose from enhanced cross-bridge availability rather than altered cytosolic Ca²⁺ signaling.

The substantial effect of small shifts in myosin ON/OFF distribution underscores strong dynamic amplification through cooperativity. Simulations with steady-state [Ca^2+^]_i_ elevation showed nearly identical steady-state forces regardless of initial ON/OFF distribution or muscle length due to cooperative activation (Fig. S5A). Without cooperativity, a ∼30 % shift in ON/OFF ratio (Fig. S6A,B) was required to match the developed force increase produced by a 4 % shift with cooperativity; while at maximal steady-state [Ca^2+^], force followed set ON/OFF distributions responding to muscle lengthening accordingly (Figs. 4L, S6A). However, lack of cooperativity lead to excessive resting tension and impaired contraction kinetics mainly due to slow relaxation (Fig. S6C-F).

In summary, modeling demonstrates that MyBP-C haploinsufficiency induces a sarcomere-driven, functionally uncoupled contractile phenotype characterized by length-dependent hypercontractility and increased calcium sensitivity without changes in calcium signaling dynamics. Cooperative interactions among sarcomeric components play a central role in amplifying MyBP-C effects on dynamic features of contraction.

## Discussion

In search of the initial cellular events preceding hypertrophic cardiomyopathy (HCM) development, this study aimed to identify the initial pre-pathological effects of a truncating mutation in myosin-binding protein C (MYBPC3-Q1061X) using patient-derived iPSC-engineered heart tissues. Elucidating the mechanisms and temporal sequence of events triggered by HCM-causing variants has been challenging, largely due to the extensive phenotypic heterogeneity observed across experimental models. A major obstacle in defining the primary, mutation-induced changes underlying pathological remodeling is that most detectable phenotypes in both model systems and patient samples reflect varying degrees of disease progression. Consequently, early molecular and functional alterations directly attributable to the causative mutation often remain obscured by secondary remodeling processes. Previous studies using MYBPC3 mutant iPSC-derived cardiomyocytes have reported that the early phenotypes are associated with secondary changes such as transcriptional remodeling (Cohn et al., 2019) and altered calcium handling (Pioner et al., 2023). In contrast, by examining tissues prior to the onset of remodeling, our results suggest that pure sarcomere driven hypercontractility precedes these downstream adaptations.

Patient-specific and genetically engineered iPSC-derived cardiac models could provide human-relevant platforms with a controllable experimental environment enabling time-resolved analysis of disease progression. However, all cardiac models originating from stem cells, including differentiated cardiac cells and physiologically more mature EHTs (Lemoine et al., 2017) present challenges related to cellular immaturity and biological as well as technical variability, thereby complicating robust qualitative and quantitative comparisons across studies (Ewoldt et al., 2025). Considering these limitations, we used iPSC-derived EHTs primarily as a controlled experimental framework to identify the initial effects of a specific gene variant preceding later stages of HCM progression. In accordance with the “two-hit” hypothesis, the onset and severity of HCM are influenced not only by pathogenic genetic variants but also by modifying genetic factors and physiological cardiac loading (Marian, 2021). To minimize conditions that might trigger remodeling, EHTs were cultured without additional hormones or growth factors and maintained at short muscle length, where tissue contraction was not significantly altered by the studied genetic variant. Importantly, all comparisons were performed between isogenic tissues generated under identical conditions, supporting the notion that the observed differences arose from the MYBPC3 mutation rather than differences in maturation state.

The differentiation protocol based on modulation of WNT signaling has been shown to promote the generation of multiple cardiac cell types (Rockel et al., 2025). Similarly, the protocol used in this study produced EHTs containing the major mesodermal cardiac cell types, including cardiomyocytes, fibroblasts, and endothelial cells, thereby mimicking aspects of *in vivo* cardiac tissue development. In the fetal human heart, these major cell types exhibit spatial and temporal shifts in their transcriptional profiles associated with emerging cardiac structures (Cui et al., 2019). Although EHTs do not recapitulate this anatomical organization, they contain populations of developing cells that resemble those in the fetal heart (Cui et al., 2019). While EHTs lack higher-order structural features such as distinct cell-type specific layers or anatomical specialization, the presence of multiple cardiac cell types enables more physiologically relevant intercellular communication, which is important for achieving functional competence (Munawar and Turnbull, 2021; Tiburcy et al., 2017). Notably, our results show that EHT cardiomyocytes express contractile protein isoforms associated with the fetal developmental stage. While isoform shifts during maturation influence multiple aspects of sarcomere function (Ahmed et al., 2022), functional properties of EHTs, including hypercontractility induced by MyBP-C deficiency and preserved Frank–Starling behavior, recapitulate key features of adult cardiac tissue.

The immediate effects of MyBP-C haploinsufficiency arise from altered well described interactions (Moss et al., 2015) between MyBP-C and other sarcomeric components that regulate myosin recruitment and cross-bridge cycling. Secondary adaptations associated with HCM-related hypercontractility include altered transcription of genes involved in Ca^2+^-signaling, membrane currents, mechanosensing, and metabolism (De Lange et al., 2023; Farrell et al., 2018), which may initially counteract primary sarcomeric dysfunction (Pioner et al., 2023). To assess these changes, we performed snRNA-seq to compare cell type–specific transcriptional signatures between MYBPC3-Q1061X EHTs and isogenic controls. We did not detect consistent transcriptional remodeling of HCM-associated gene patterns in cardiomyocytes or other cardiac cell types. In line with this, we observed no evidence of hypertrophy at the tissue or cellular level, nor HCM-associated alterations in metabolism, respiration, or calcium signaling. While subtle changes below the detection threshold cannot be excluded, the absence of consistent alterations across transcriptional, metabolic, and functional readouts strongly supports the conclusion that overt remodeling is absent under these conditions. Together, these findings indicate that the hypercontractile phenotype observed here represents a primary consequence of MyBP-C haploinsufficiency rather than a secondary adaptation and likely precedes the onset of HCM remodeling process.

The hypercontractile phenotype observed in the present study is consistent with the fundamental relationship between force generation and cross-bridge recruitment (Moss et al., 2004), whereby increased force arises from greater availability of myosin heads for actin interaction (Spudich, 2015). Reduced MyBP-C enhances the probability of cross-bridge formation by destabilizing the super-relaxed (SRX) state and release of myosin head tethering (Spudich et al., 2024). Accordingly, the truncating *MYBPC3* variant induces hypercontractility characterized by increased force generation and accelerated force development (Korte et al., 2003). In this study, we further show that this phenotype, including increased force, contraction kinetics, and ATP turnover, is dependent on muscle length. In principle, both length-dependent activation (LDA) (Campbell et al., 2018; Ma et al., 2021), and reduced MyBP-C levels (McNamara et al., 2017; McNamara et al., 2016) increase recruitment of OFF-state myosins, suggesting that these mechanisms could act additively, provided that myosin availability is not limiting. However, the majority of studies examining the impact of MyBP-C deficiency on LDA suggest that the absence of MyBP-C suppresses length-dependent force generation (Hanft et al., 2021; Mamidi et al., 2014; Najafi et al., 2016). Since hypercontractility due to MyBP-C deficiency is followed by secondary adaptations that suppress hypercontractility (De Lange et al., 2023; Pioner et al., 2023), variability across studies likely reflects differences in the extent of remodeling. Our experiments with hypercontractile EHTs showing no signs of secondary remodeling demonstrated that the central characteristics of hypercontractility, namely increased force development speed and augmented force, are more pronounced at longer muscle lengths. While a contribution of EHT immaturity to this phenotype cannot be entirely ruled out, our modeling suggests that, within the primary assumptions of the model, the adult sarcomere recapitulates the same phenomenon. Notably, the modeling further suggests that hypercontractility fine-tunes sarcomeres and primarily affects the dynamic features of contraction, i.e., the response to transient elevations of [Ca²⁺]ᵢ. This highlights that methodological differences between intact and permeabilized preparations, as well as variations in experimental muscle length, may contribute to variability across studies.

Marked augmentation of force development due to disrupted physiological control, such as hypercontractility resulting from MyBP-C deficiency, may impair normal cardiac function. In our experiments, increasing beating frequency at increased muscle length led to elevated baseline tension in MYBPC3-Q1061X EHTs, a feature that corresponds to diastolic dysfunction in the intact heart. Although extrapolation from EHTs to *in vivo* physiology is not straightforward, hypercontractility is likely to have the greatest impact under conditions of high ventricular volume and low β-adrenergic drive, as PKA-dependent phosphorylation can partially counteract the effects of MyBP-C haploinsufficiency (Nagayama et al., 2007). According to our results, hypercontractility is more prominent at longer muscle lengths, suggesting that this phenotype does not simply augment the contractility of cardiac muscle but also modifies the FS-response of the heart. The consequences of hypercontractility may be more pronounced in isolated systems that lack systemic regulation of relaxation kinetics, but our findings nonetheless qualitatively recapitulate calcium sensitization-related diastolic dysfunction observed in heterozygous knock-in mice (Fraysse et al., 2012), the high prevalence of diastolic dysfunction in HCM patients (Dereli Bulut et al., 2026), and early pre-hypertrophic dysfunction in carriers of the MYBPC3-Q1061X variant (Poutanen et al., 2006).

To investigate the biophysical origin of the hypercontractile phenotype, we employed a thermodynamically consistent cross-bridge model which incorporates sarcomeric cooperativity. Model parameters were constrained by previously established experimental measurements of sarcomeric kinetics, allowing us to test whether shifts in myosin state equilibrium were sufficient to reproduce the observed phenotype. Notably, the qualitative behavior of the model was robust across physiologically plausible parameter ranges, indicating that the conclusions were not dependent on fine-tuned parameter selection. The simulations indicated that a relatively small (∼4%) shift in the myosin ON/OFF equilibrium, corresponding to the SRX/DRX transition, was sufficient to recapitulate both the magnitude of hypercontractility and its length dependence. This amplification is consistent with cooperative regulation within the sarcomere (Alamo et al., 2017; Moss et al., 2015; Nelson et al., 2023). In the absence of cooperativity, substantially larger shifts in the ON/OFF equilibrium are required to achieve similar force increases, leading to non-physiological levels of diastolic tension and markedly slower contraction kinetics. These results indicate that sarcomeric cooperativity amplifies modest changes in myosin state into substantial functional effects while preserving physiological contraction dynamics. Furthermore, simulations suggest that small shifts in myosin ON/OFF ratio would have limited functional consequences in the absence of cooperativity.

Our model incorporates nearest-neighbor interactions among regulatory units and cross-bridges. While cross-bridge cooperativity following MyBP-C–dependent shifts in ON/OFF ratio increases recruitment of force-generating units, additional interactions are required to fully explain the hypercontractile phenotype (Kalda and Vendelin, 2020). Because Ca^2+^ binding to the thin filament does not reach equilibrium during the cytosolic Ca^2+^ transient, force production depends on both the amplitude and duration of Ca^2+^ signals (Moss et al., 2015), whereas thin filament activation kinetics ultimately limit the rate of force development (Regnier et al., 2004). Consistent with our experimental observations, thin filament cooperativity allows increased force generation without corresponding increases in Ca^2+^ binding (Campbell et al., 2010). Accordingly, when Ca^2+^ binding to troponin C is unchanged, cytosolic Ca^2+^ buffering and transient waveforms remain unaffected. This indicates that hypercontractility can arise from intrinsic sarcomeric biophysical modulation without alterations in cytosolic Ca^2+^ signaling or in downstream pathways. Importantly, the model reproduces both the magnitude of hypercontractility and its length dependence using a single mechanistic perturbation, supporting the conclusion that altered myosin state equilibrium is sufficient to explain the observed phenotypes.

Under the controlled conditions used in this study, the biophysical consequences of reduced MyBP-C did not trigger detectable secondary adaptations. This suggests that the mutation itself does not directly initiate EC-coupling remodeling which contributes to the phenotype but rather leads to adaptive responses which arise from the altered mechanical and energetic demands imposed by hypercontractility over time. Although such conditions cannot fully replicate the *in vivo* environment, they highlight a central principle of HCM pathogenesis, disease progression reflects the interaction between genetic predisposition and additional physiological stress (Maron et al., 2022). Mechanistically, two immediate consequences of hypercontractility, elevated energy consumption and altered mechanical stress, are likely to initiate downstream pathways driving hypertrophic remodeling, consistent with proposed mechanisms of HCM (Ranjbarvaziri et al., 2021; Sequeira et al., 2023) and general forms of hypertrophy (Lyon et al., 2015; Ritterhoff and Tian, 2023; Tuomainen and Tavi, 2017). While our data supports sarcomere-driven hypercontractility as an early event preceding remodeling, establishing direct causal links to downstream pathological changes will require further time-resolved studies under conditions favoring disease progression.

In conclusion, we investigated the earliest cellular effects of the truncating MYBPC3-Q1061X mutation using patient-derived iPSC-engineered heart tissues. Under conditions that minimize external remodeling stimuli, we observed no evidence of transcriptional, metabolic or hypertrophic remodeling, whereas a robust hypercontractile phenotype was preserved. Computational modeling supports the notion that small shifts in myosin ON/OFF states, amplified by sarcomeric cooperativity, are sufficient to explain this phenotype without requiring alterations in calcium signaling. These findings indicate that hypercontractility is the primary effect of MyBP-C haploinsufficiency that precedes secondary remodeling processes. As pharmacological agents targeting myosin activity have emerged as promising therapies for HCM, modulation of myosin state equilibrium or sarcomeric cooperativity may offer opportunities for early intervention, potentially preventing disease progression rather than reversing established pathology.

## Methods

### Induced pluripotent stem cell generation and culture

Induced pluripotent stem cells were produced from fibroblasts acquired from skin biopsies of two patients with hypertrophic cardiomyopathy heterozygous for the truncating MYBPC3-Q1061X variant after informed consent and approval from the committee on Research Ethics of Northern Savo Hospital district (license no 64/2014). Fibroblasts were reprogrammed with CytoTune™-iPS 2.0 Sendai Reprogramming Kit (Invitrogen), clonal iPSC colonies were selected and the iPSC phenotype was confirmed as previously described (Oksanen et al., 2017). Cells were maintained on Matrigel (Corning) coated cell culture dishes in E8 medium (Gibco) supplemented with 0.5 % penicillin-streptomycin (P/S) (Gibco) under standard cell culture conditions. Cells were passaged twice a week by dissociation with EDTA.

### Genome editing

The diseased Q1061X variant in patient-derived iPSCs was repaired using CRISPR-Cas9 nickase-based genome editing (Ran et al., 2013). Relative to the NCBI dbSNP entry for the variant (rs397516005), PAM sites 8 base pairs downstream on the opposite strand and 50 base pairs upstream on forward strand were targeted by cloning the 20 base pair target sequences into pSpCas9n(BB)-2A-Puro (PX462) V2.0 plasmid (a gift from Feng Zhang, Addgene plasmid #62987) according to the provided protocol. DNA oligo sequences for the cloning were (5’-3’) CACCGACGCTGGTGCTGTAGGTTGT and AAACACAACCTACAGCACCAGCGTC for the opposite strand target, and CACCGTCTCAATGCGCACCGTCACC and AAACGGTGACGGTGCGCATTGAGAC for the forward strand target. The opposite strand target contained the variant nucleotide, and thus the sequence cloned into the plasmid contained the nucleotide corresponding to the HCM-associated variant. iPSCs were transfected with the plasmids together with DNA template for homology-directed repair (150 base pair oligo on forward strand, corrected base on position 76) using a Neon™ Electroporation System (Invitrogen) according to the manufacturer instructions. After plating the transfected cells, puromycin selection was performed and surviving clonal cell colonies were selected and further expanded. CRISPR-Cas9 clones were screened for the successful editing using restriction enzyme analysis for the Q1061X variant as previously described (Jaaskelainen et al., 2002) and confirmed by Sanger sequencing of the region surrounding the variant nucleotide.

### iPSC differentiation into cardiomyocytes

For cardiomyocyte differentiation, a previously described method utilizing WNT pathway modulation with synthetic small molecules was used (Burridge et al., 2015; Lian et al., 2012). EDTA-dissociated iPSCs were plated on Matrigel coated 24-well-plate wells and differentiation was initiated at 90-100 % confluency, 2-3 days after plating. First, the E8 medium was changed to CDM3 medium (RPMI 1640 (Gibco) with 0.1 % bovine serum albumin (Sigma), 0.033 mM ascorbic acid (Sigma) and 1 % P/S) supplemented with 9 µM CHIR99021 (Tocris). Twenty-four hours later, medium was changed to CDM3 without supplementation and cells were cultured in this solution for 48 h. Next, the cultures were kept in CDM3 supplemented with 2 µM Wnt-C59 (Tocris) for 48 h after which CDM3 without supplementation was used until the cells were used for experiments. Dissociation of differentiated cells into single cells was performed as previously (Naumenko et al., 2024) with a buffer solution (NaCl 100 mM, KH_2_PO_4_ 1.2 mM, MgSO_4_ 4 mM, HEPES 10 mM, Taurine 50 mM, Glucose 20 mM, KCl 10 mM) containing 2 mg/mL type II collagenase (Worthington) and 2 mg/mL pancreatin (Sigma-Aldrich).

### Flow cytometry

The proportion of cardiomyocytes in differentiation cultures was determined by flow cytometric analysis of cardiac Troponin T as previously (Naumenko et al., 2024). Dissociated cells from differentiation cultures were fixed with 2% paraformaldehyde (in PBS) for 10 min and permeabilized with 5 % saponin in PBS for 15 min. Cells were incubated with Anti-Cardiac Troponin T-FITC antibody (Miltenyi Biotec GmbH) for 30 min and the fluorescence signal was analyzed using a CytoFLEX Flow Cytometer (Beckman Coulter). Non-stained samples were used to determine the background fluorescence and based on this, gating for Troponin T positive cells was performed.

### Engineered heart tissue generation and culture

The protocol for Engineered heart tissue (EHT) generation was adapted from a previously published method (Breckwoldt et al., 2017). PDMS silicone racks (EHT Technologies) for EHT culture, designed to be used as an array of four tissues, were cut into single units to allow for the isometric force recording with our custom-made setup. Agarose molds were prepared in 24-well-plates using Teflon spacers (EHT Technologies) and boiled solution of 2 % agarose in PBS. After agarose solidification, Teflon spacers were removed and single-unit PDMS racks were positioned at the center of the formed agarose molds. For tissue casting, 50 µL of solution containing 10 mg/mL fibrinogen (Sigma), 10 µL Matrigel, 2 µg/mL aprotinin (Sigma) and 10 µM Rho kinase inhibitor Y-27632 (Selleck Chemicals) in DMEM was combined with 47 µL of solution containing one million cells and 2 % P/S in DMEM. The resulting solution was mixed with 3 µL of 1000 U/mL thrombin (Sigma) and quickly pipetted into the agarose mold containing the PDMS rack. The final concentration of each component in the resulting solution was 10^7^ cells/mL, 5 mg/mL fibrinogen, 3 U/mL thrombin, 100 µL/mL Matrigel, 1 µg/mL aprotinin, 5 µM Y-27632 and 1 % P/S in DMEM. The plate was placed in the cell incubator for 80 min after which 500 µL DMEM (1 % P/S) was added and after a further 10 min incubation, PDMS racks with solidified EHTs attached to the rack posts were carefully moved into new 24-well-plate wells with EHT medium (DMEM with 10 % horse serum (Gibco), 33 µg/ml aprotinin, 10 µg/ml insulin (Thermo Scientific) and 1 % P/S). EHTs were maintained in 24-well-plates and five days before they were used in force recordings or sample preparation, they were placed in constant 1 Hz electrical stimulation using a C-Pace Culture Stimulator System (Ionoptix) to minimize the effects of slight variability in the spontaneous contraction frequency. For the electrical stimulation, a PDMS rack with EHT was placed between the carbon electrode plates of C-Dish electrodes designed for 35 mm dishes (Ionoptix) and the electrode plate was placed on a 6-well-plate containing enough EHT medium in each well to submerge the EHT and the ends of the carbon electrode plates.

### EHT cell dissociation and culture

For 2D experiments of cells cultured in EHTs, cells were dissociated with type II collagenase and pancreatin as from the differentiation cultures before EHT generation. Dissociated cells were re-suspended into 2D culture medium (DMEM with 10 % fetal bovine serum (Gibco) and 1 % P/S) supplemented with 10 µM Y-27632 and plated on mouse laminin (Sigma) coated glass coverslips or XF96 cell plates (Agilent Technologies, Santa Clara, CA, USA) at a density of 65000 cells/cm^2^ or 470000 cells/cm^2^, respectively. The next day, the medium was changed to 2D culture medium without Y-27632. Cells were assayed two days after plating.

### Western blot

Total protein from snap-frozen EHTs stored in –70 °C was isolated as previously described for mouse heart tissue (Mutikainen et al., 2016). Twenty-five µg of protein was separated with SDS-PAGE and transferred to 0.2 µm nitrocellulose membrane. The membrane was incubated with rabbit anti-MYBPC3 (Abcam) antibody diluted 1:100 in 5 % BSA in TBST at 4 °C overnight. For secondary HRP-conjugated anti-rabbit (Cell Signaling Technologies) antibody, 1:10000 dilution in 2 % BSA in TBST at RT for one hour was used. The blot was imaged using GelDocTM MP imaging system (Bio-Rad Laboratories Inc.) and quantified using ImageJ software. MYBPC3 protein band signal was normalized to total protein transferred surrounding the band as determined with Ponceau S staining (Sigma-Aldrich).

### Contraction force measurements

Direct isometric force measurement was performed in a custom-made setup consisting of a perfusion chamber, platinum wires for electrical stimulation, a micromanipulator for EHT positioning, an isometric force transducer (HSE-HA F10 type 375) and a remote-controlled motorized lever on a confocal inverted microscope (FluoView 1000; Olympus, Tokyo, Japan). Experiments were performed under constant perfusion of carbogenated DMEM (37 °C, pH 7.4) and constant 1.5 Hz electrical stimulation. Before the force recording, a PDMS rack with EHT was attached to the micromanipulator arm and the EHT positioned into the chamber and perfused without stretching for 10 min while electrical pacing. The PDMS rack was positioned with the micromanipulator so that the arms of the force transducer and motorized lever were on the opposite posts of the PDMS rack right above the EHT. The response of EHTs to increase in the muscle length was assessed from slack length to the length of maximal force development with stepwise stretching at the speed of 60 µm/min with the remotely controlled motor lever. From the recorded force trace, the length at which the EHT produced half of the maximum twitch force was evaluated and length adjusted. The EHT was perfused at the length of half-maximal force for 10 min and then exposed to a pacing frequency protocol, where the EHT was paced at 0.5, 1, 2 and 3 Hz for 20 s each. After the experiment, the EHT was either snap frozen with liquid N_2_ and stored in –70 °C or dissociated into single cells.

### Single-nuclei RNA sequencing and data analysis

Nuclei extraction from frozen EHTs and droplet-based single-nuclei RNA library preparation using the 10X Genomics Chromium platform (10X Genomics) were performed as previously described for adult human cardiac tissue (Linna-Kuosmanen et al., 2024). Indexed cDNA libraries were pooled and sequenced using a NovaSeq 6000 S2 system (Illumina). Single-nuclei gene counts were obtained using the Cell Ranger software (10X Genomics) with default parameters and the provided GRCh38 reference file. Next, Velocyto (v.0.17.17) (La Manno et al., 2018) was run while masking repeat regions, as demonstrated in their tutorial. The output was used to calculate the fractions of unspliced reads per barcode. This information was then used to run QClus (v.0.2.0) (Schmauch et al., 2025), which automatically ran Scrublet (v.0.2.3) (Wolock et al., 2019) as part of its pipeline. In addition to QClus, CellBender (v.0.2.2) (Fleming et al., 2023) was run independently. To decontaminate the counts and remove droplets, the barcodes filtered out by QClus were removed from the CellBender output. Further data processing to achieve UMAP plots and Leiden clustering was performed using Python with the functions of the Scanpy toolkit (v.1.11.1( (Wolf et al., 2018) and Harmony batch correction (Korsunsky et al., 2019) followed by cell type annotation as previously described (Linna-Kuosmanen et al., 2024). Previously published snRNA-seq data from the left ventricles of human HCM patients (Chaffin et al., 2022) were processed as EHT data. For the integration of EHT and human snRNA-seq data, subsets of common cell types (cardiomyocytes, fibroblasts, and endothelial cells) from both datasets were combined and integrated by running the Harmony batch correction first at the sample level and then at the dataset level. For gene expression changes in snRNA-seq data, single-nuclei count matrices were pseudobulked at the cell type level, and the *DESeq2* package (*v1.50.2*) in R 4.5.2 software (R Core Team, 2025) was used for differential expression analysis. QIAGEN IPA (QIAGEN Inc., https://digitalinsights.qiagen.com/IPA) was used for pathway analysis as previously described (Linna-Kuosmanen et al., 2024).

### Metabolite analysis with LC-MS/MS

Non-targeted metabolite profiling was performed at the Biocenter Kuopio LC-MS Metabolomics Center at the University of Eastern Finland. EHTs stored in –70 °C were placed in 40 µL of deionized water in 1.5 mL Eppendorf tubes on ice and sonicated for 5 min. Next, 240 µL of acetonitrile with 0.1 % formic acid was added while mixing with vortex shaker, after which samples were further sonicated for 5 min. Samples were filtered with 0.2 µm polytetrafluoroethylene syringe filters (Pall). Two microliters of samples were separated with either reverse phase (RP, Zorbax Eclipse XDB-C18, 2.1 μm 100 mm, 1.8 mm column, Agilent Technologies, Palo Alto, CA, USA) or hydrophilic interaction (HILIC, Acquity UPLC BEH Amide 1.7 μm, 2.1 × 100 mm column, Waters, Ireland) chromatography columns using an ultra-high performance liquid chromatography (UHPLC) system (Vanquish Flex UHPLC system, Thermo Scientific, Bremen, Germany). The sample order was randomized, and after every 10 samples quality control, prepared by pooling all samples, was injected. The raw data was processed with MS-DIAL software (version 4.9) and features were identified using a local metabolite library maintained by the University of Eastern Finland Metabolomics Center, and publicly available mass spectrometry databases (METLIN (https://metlin.scripps.edu), MassBank of North America (MoNA, https://mona.fiehnlab.ucdavis.edu), Human Metabolome Database (HMDB, www.hmdb.ca), and LIPID MAPS (https://www.lipidmaps.org). Total number of detected molecular features after initial filtering was 5454 distributed among different chromatography and data acquisition modes as follows: RP with ESI+ (RPpos) 2976, RP with ESI-(RPneg) 1149, HILIC with ESI+ (HILICpos) 733, HILIC with ESI-(HILICneg) 596. For the final statistical analysis, only features that were automatically annotated in MS-DIAL software by matching at least exact metabolite mass and MS/MS spectrum were chosen, leading to the final number of 452 metabolites (RPpos 237, RPneg 50, HILICpos 110, HILICneg 55). The most frequent metabolite types were phosphatidylcholines (PC), phosphatidylethanolamines (PE), diglycerides (DG) and acylcarnitines (ACAR) with 112, 70, 38 and 24 molecules, respectively.

### Analysis with Seahorse extracellular flux analyzer

One day after the dissociation and plating of EHT cells on XF96 cell plates, cells were assayed using a Seahorse XFe96 analyzer (Agilent Technologies, Ca, Santa Clara, USA). Cells were washed once with XF Base Medium (Agilent Technologies, CA, Santa Clara, USA) (pH 7.4) and 180 µL of assay medium (XF Base Medium supplemented with 2 mM GlutaMAX, 4.5 g/L glucose and 0.11 g/L sodium pyruvate (Sigma-Aldrich, MO, St. Louis, USA)) was added to the wells. A mitochondrial stress test was performed by sequential injections of oligomycin A, carbonyl cyanide 4-(trifluoromethoxy)phenylhydrazone (FCCP) and antimycin A to final concentrations of 1, 1.5 and 5 µM, respectively. Oxygen consumption and extracellular acidification rates from each well were normalized to total protein in the well assessed using the Bradford method.

### Action potential recordings

Coverslips with attached cells were transferred to the recording chamber of the microscope and perfused with preheated DMEM solution (37C°, pH=7.4, bubbled with 95% O2 and 5% CO2). For action potential (APs) recordings patch-clamp amplifier Axopatch 200B in combination with a Digidata 1440A and Clampex 10 software (Molecular Devices Inc., Sunnyvale, CA, USA) were used as previously described (Naumenko et al., 2021). An Ag/AgCl half-cell electrode (World Precision Instruments Inc., Sarasota, FL, USA) connected to the bath via an agar bridge was used as the ground electrode. Patch pipettes were pulled from borosilicate glass capillary tubing (ID 0.86 mm, Harvard Apparatus, Edenbridge, UK) with a micropipette puller (Sutter P-97, Sutter Instrument Company, Novato, CA, USA) and fire-polished. The patch pipette resistances were 4–6 MΩ when filled with the pipette solution containing (in mmol/L): K-aspartate 120, KCl 8, NaCl 7, MgCl_2_ 1, Na_2_-phosphocreatine 2, Mg-ATP 5, Na-GTP 0.3,and HEPES 10 (pH 7.20 adjusted by KOH). APs were recorded using the patch-clamp whole-cell configuration in current-clamp mode (I=0). Evoked APs were elicited by brief (1 ms) suprathreshold current pulse. Analysis of AP recordings was performed with ClampFit 10 (Molecular Devices, Sunnyvale, CA, USA).

### Confocal calcium imaging

Calcium imaging was performed as previously described (Koivumäki et al., 2018). Coverslips with attached cells were transferred to the Dulbecco’s Modified Eagle Medium plus GlutaMAX I (DMEM, Life Technologies, Carlsbad, CA, USA) with Fluo-4-acetoxymethyl (AM)-ester (5 μM, Invitrogen, Carlsbad, CA, USA) for 30 min at 37 °C. The cells were then placed in a recording chamber (Cell MicroControls, Norfolk, VA, USA, flow rate approx. 1-2 ml/min), where they were continuously perfused with preheated DMEM and equilibrated with carbogen gas. To allow de-esterification of the dye the cells were perfused for 20 min before the experiment. All experiments were carried out at 37°C sustained by a temperature controller (TC2BIP, Cell MicroControls, Norfolk, VA, USA). Only well-attached cells showing regular spontaneous contraction were used for recordings. Measurements were performed with a confocal inverted microscope (FluoView 1000; Olympus, Tokyo, Japan). To measure myocyte calcium signals the cells were excited at 488 nm and the emitted light was collected at 500-600 nm through a 60X objective lens in line-scan mode. To excite the cells, myocytes were paced with 1-ms square voltage pulses 50% over the excitation threshold through two platinum wires located on both sides of the chamber. In some experiments caffeine (10 mM, Sigma) was applied directly to the studied area with a local perfusion manifold (Cell MicroControls, USA). Fluo-4 fluorescence intensity is expressed as an F/F_0_-ratio, where F is the background-subtracted fluorescence intensity and F_0_ is the background-subtracted minimum fluorescence value measured from each cell at rest. The images were analyzed using FluoView and ImageJ (imagej.nih.gov/ij/) softwares.

### L-type Ca^2+^ current recordings

Patch-clamp experiments were performed in the same recording chamber at 37°C, and the cells were perfused with normal Tyrod solution (bubbled with 100% O_2_). Whole-cell voltage-clamp (Axopatch 200B, Digidata 1440A, Molecular Devices Inc., USA) was used for the L-type Ca^2+^ current. Patch electrodes (Harvard Apparatus, United Kingdom) were pulled and fire-polished with Sutter P-97 (Sutter Instrument Company, Novato, CA). Patch electrodes for Ca^2+^-current measurements had resistances of 3-4 MΩ when filled with pipette solution. The cell capacitance and series resistance were electronically compensated. The cells with an unstable or high access resistance were discarded. Under voltage clamp control cells were held at -80 mV. Membrane capacitance and resistance were estimated in response to a 5mV pulse. The current amplitudes were normalized by cell capacitance. Recordings were carried out at a sampling rate of 10 kHz, and a low-pass Bessel filter at 5 kHz was used. L-type Ca^2+^ current recordings protocol was previously described (Xu et al., 2011). The cells were perfused with Tyrode solution containing (in mM): 130 NaCl, 5.4 KCl, 1 CaCl_2_, 1 MgCl_2_, 0.3 Na_2_HPO_4_, 10 HEPES, and 5.5 glucose, pH 7.4 with NaOH. The internal solution contained (in mM): 110 CsOH, 90 aspartic acid, 20 CsCl, 10 tetraethylammonium chloride (TEA chloride), 10 HEPES, 10 EGTA, 5 Mg-ATP_2_, 5 Na_2_-creatine phosphate, 0.4 GTP-Tris, 0.1 leupeptin (pH 7.2 with CsOH). After establishing of the whole-cell configuration, the bath solution was switched to recording solution (in mM): 125 N-methyl-glucamine, 5 4-aminopyridine, 20 TEA chloride, 2 CaCl_2_, 2 MgCl_2_, 10 glucose, 10 HEPES (pH 7.4 with HCl). After an initial 1-sec prepulse at -40 mV, Ca^2+^currents were elicited using 200-ms voltage steps from -30 to +50 mV in 10-mV increments. Voltage-dependence of inactivation was assessed by holding cells at various potentials from - 40 to +10 mV for 2 sec followed by a 100-ms test pulse to +10 mV. Electrophysiological data was analyzed using ClampFit 10.7 software.

### Modeling

The mechanical model used here is a thermodynamically consistent cross-bridge model based on the Huxley-type framework, incorporating cooperativity through the movement of tropomyosin (Kalda et al., 2015; Kalda and Vendelin, 2020). Using the standardized Ca^2+^ - transient as an input for the model, the binding of calcium or a cross-bridge induces the displacement of tropomyosin, which in turn alters the free energy landscape of the cross-bridge ensemble. Simulations were performed on infinitely long chains of cross-bridges (XBs) and regulatory units (RUs). The model tracks only a single XB, under the assumption that the neighboring XBs share the same distribution among their states. Cooperativity is simulated through nearest-neighbor interactions among RUs and XBs: RU-RU, XB-RU, and XB-XB interactions (Razumova et al., 2000). Among these interactions, the RU-RU and XB-XB interactions are mediated by changes in the free energy profile of the actomyosin reaction, and the corresponding parameters were optimized when the data were fitted. The XB-RU interaction is derived by assuming elastic deformation of tropomyosin. To account for the OFF state of myosin heads, two additional states were introduced: an OFF (super-relaxed, SRX) state without calcium bound and an OFF state with calcium bound to troponin (Fig. 4A). In the simulations, myosin heads in the OFF (SRX) state were in a super-relaxed conformation that precluded actin binding, whereas ON (disordered-relaxed, DRX) state myosin heads are available to bind actin and form force-generating cross-bridges. The transition between these states is governed by thermodynamic equilibrium and was restricted to the unbound myosin pool; thus, actin-bound myosin heads were not allowed to transition directly to the OFF state. For adult cardiomyocytes, unbound myosins were assumed to be equally distributed between ON and OFF states. Reference model parameters were then determined by fitting to isometric stress data from rat trabeculae (Janssen and Hunter, 1995) (Fig. S7A,B) and to the linear relationship between oxygen consumption and the stress–strain area (Hisano and Cooper, 1987) (Fig. S7C,D). The optimization was conducted in two stages. First, a range of values for free energy parameters describing the deformation of tropomyosin and transitions between different myosin states were specified. Then, for all the possible combinations of those parameters, the corresponding rate constants describing cross-bridge binding and interaction with tropomyosin were optimized using least squares minimization. This fitting procedure was performed separately for two different sarcomere lengths (1.9 and 2.1 µm), resulting in distinct parameter sets for each case. The parameter sets that resulted in a satisfactory fit with minimal residual error were used in additional simulations, where the proportion of myosin heads in the OFF state was varied from 50 to 100% in 0.5% increments.

### Data analysis and statistics

Results are presented as mean values and standard error of the mean (SEM) is used to describe data variance, unless stated otherwise. A two-tailed Student’s *t*-test was used for statistical testing of data with two experimental groups and one-way ANOVA with Bonferroni *post hoc* test for data with four experimental groups. For the metabolomics data, adjusted α level for statistical significance of multiple testing was acquired by dividing value 0.05 with the number of principal components required to explain 95 % of the variance within data. For the analysis of EHT Frank-Starling and pacing frequency responses, a linear mixed effect model using *lmer* function of *lme4* package (*v2.0-1*) (Bates et al., 2015) followed by statistical testing of model estimates using functions of *emmeans* package (*v2.0.2*) (https://rvlenth.github.io/emmeans/) in R software were used. In all tests, *P* < 0.05 was considered statistically significant.

## Author contributions

**Conception and design of the experiments:** P.T., T.T., N.N., M.V., S.L.-K. **Collection, analysis and interpretation of data:** P.T., T.T., S.L.-K., M.S, M.R., J.T.K., E.S.,K.G., N.N. **Drafting the article and reviewing:** P.T., M.S., M.R., M.V., S.L.-K., T.T., R.L., J.T.K., N.N., J.K. **Resource contribution:** P.T., S.L.-K., M.K., K.G. **Materials provision:** J.K., P.T.

## Funding

Bristol Myers Squibb (ISR) #CV027-052 (P.T., J.K); Sigrid Juselius Foundation # 250237 (P.T.), #250133 (S.L.-K.); Research Council of Finland # 325510 (P.T.), #342074 (S.L-K.)

## Conflict of Interest

None

## Acknowledgments

The authors thank Anne Karppinen and Jalmari Laurila for their help with cell culture and sample preparation.

## Supplementary Information

**Supplementary Figure 1.**
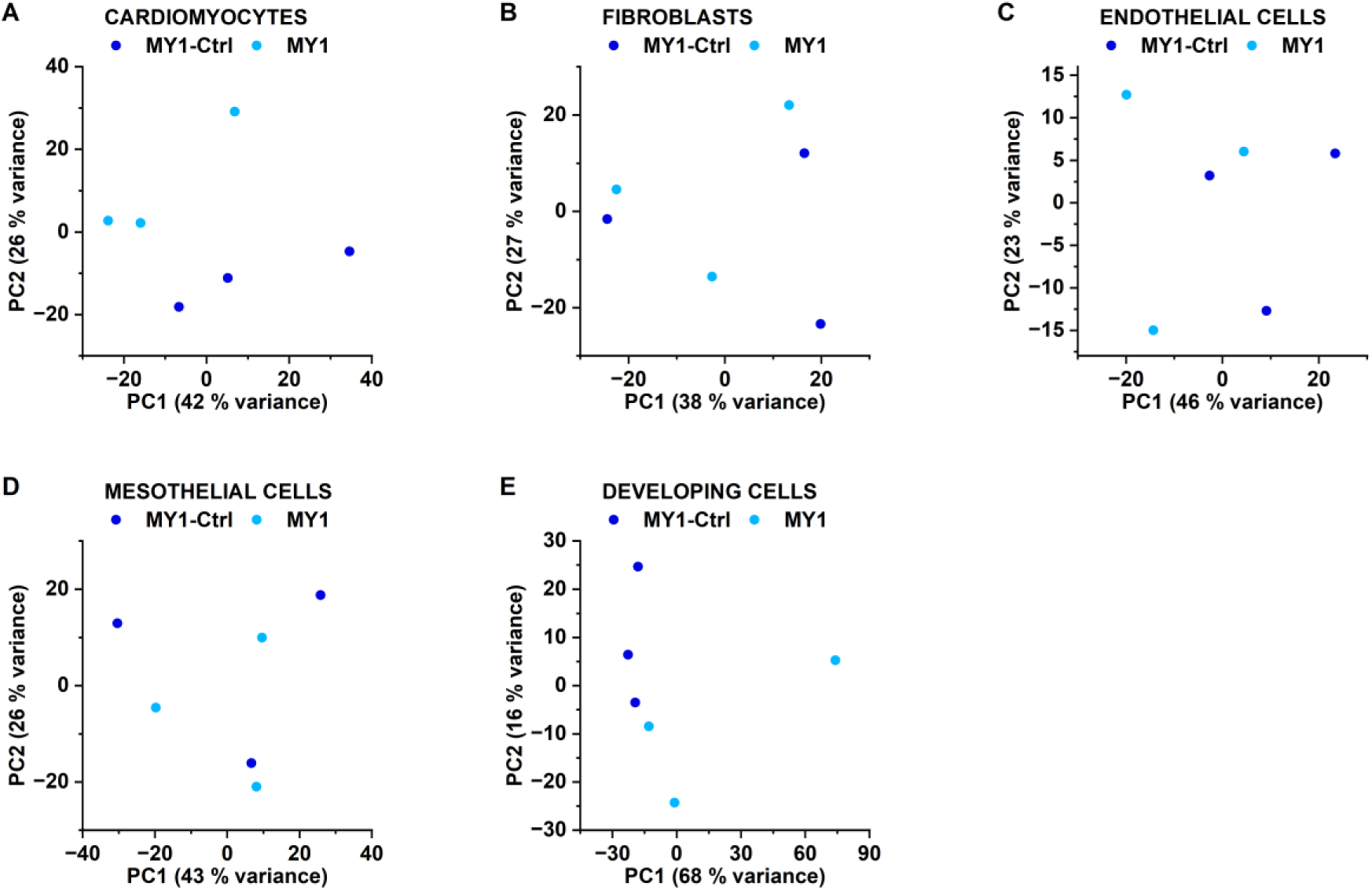
Principal component analysis of cell-type specific gene expression. **A)-E)** PCA performed with *DESeq2* software on cell-type specific pseudobulk gene expression data for A) cardiomyocyte, B) fibroblast, C) endothelial cell, D) mesothelial cell and E) developing cell populations.

**Supplementary Figure 2.**
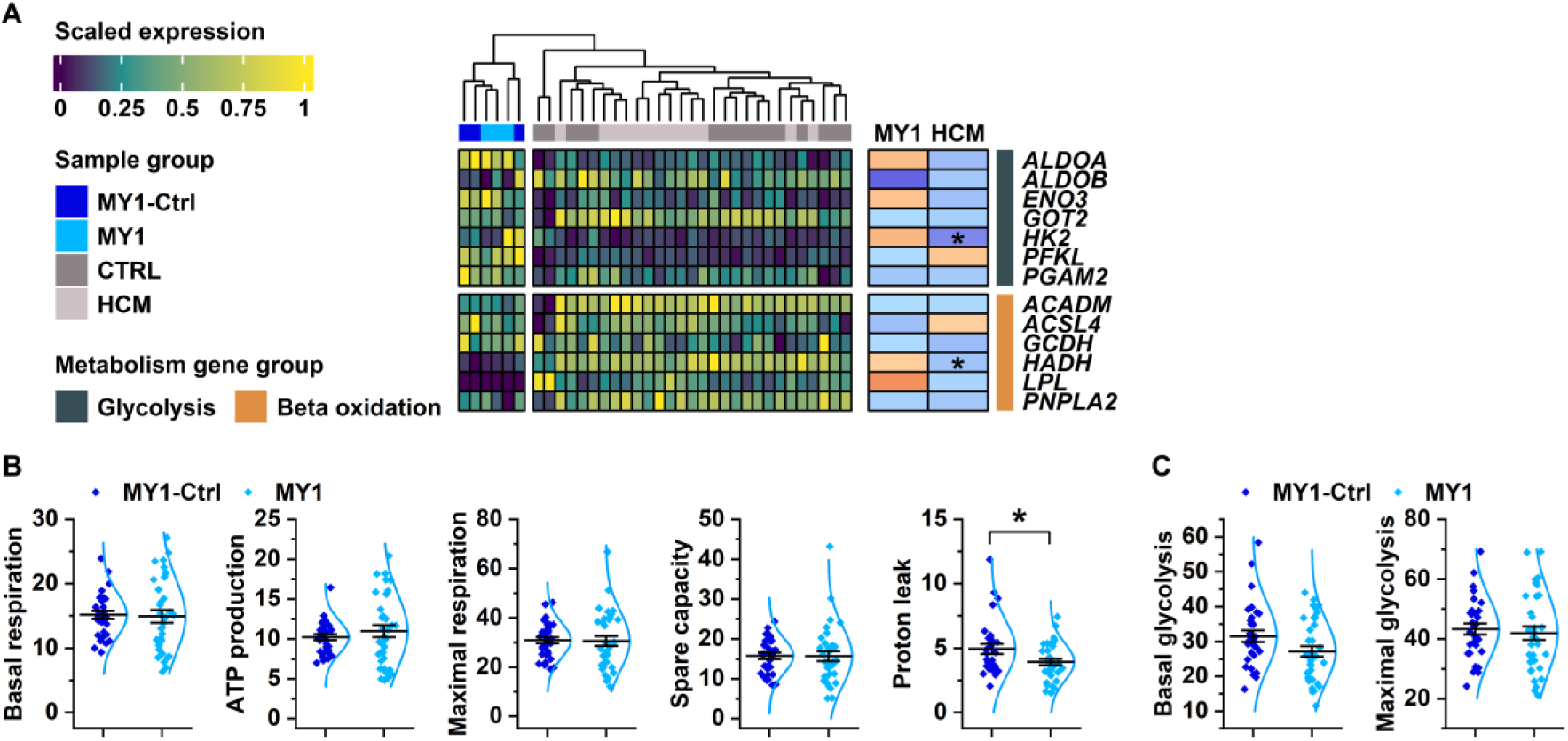
Energy metabolic phenotype of MYBPC3 Q1061X carrying cardiomyocytes. **A)** Expression level and changes of glycolysis and beta oxidation related genes not shown in the heatmap in Figure 2E. **B)-C)** Individual OCR (B) and ECAR (C) parameters assessed from mitochondrial stress test. n(MY1-Ctrl)=22, n(MY1)=24. Student’s *t*-test *P* value: *, *P* < 0.05.

**Supplementary Figure 3.**
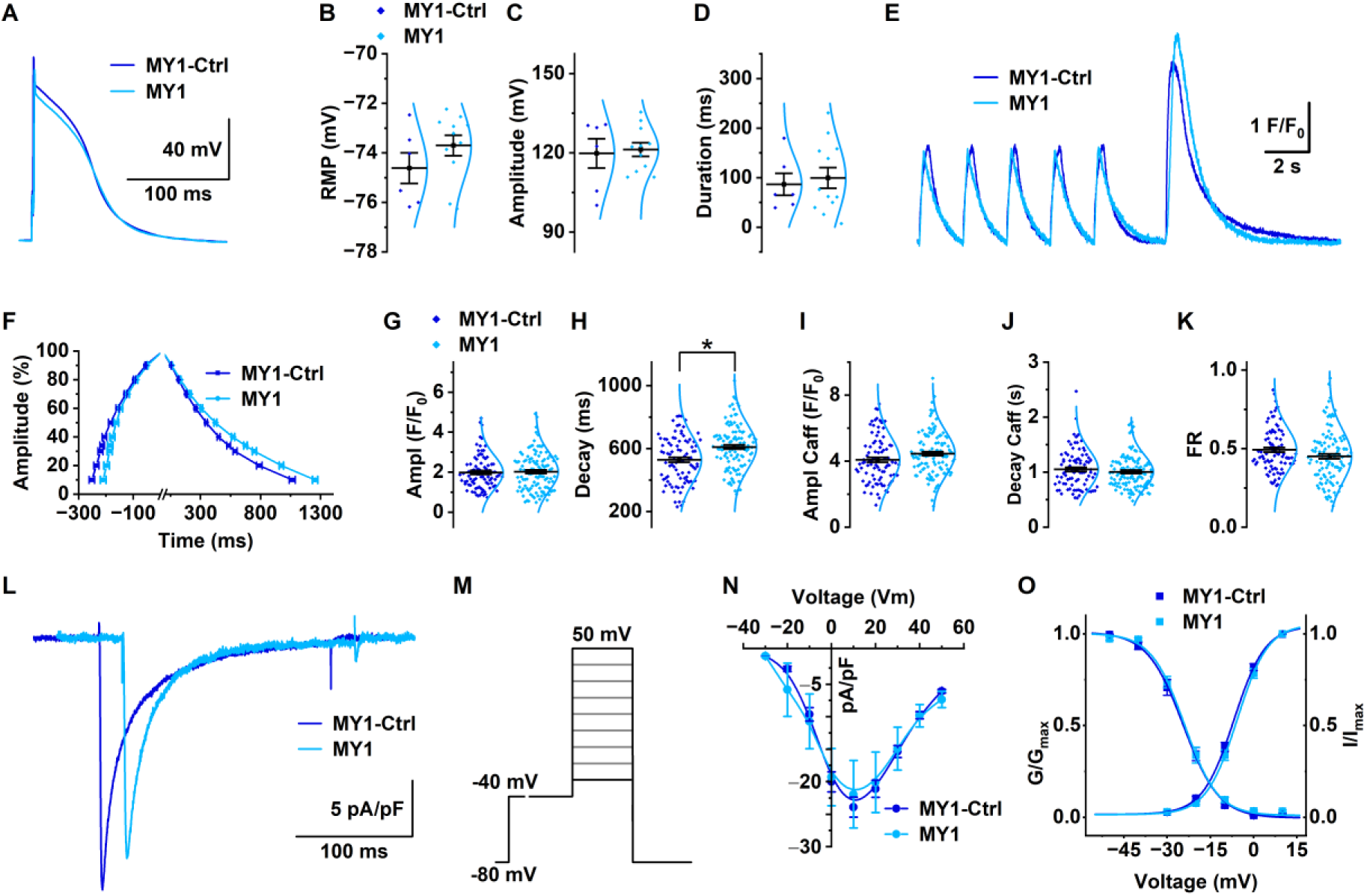
Calcium transient and L-type calcium current in cardiomyocytes isolated from EHTs. **A)-D)** Representative traces (A), resting membrane potential (B), amplitude (C) and duration (D) of cardiomyocyte action potential. In B, C and D n(MY1-Ctrl)=6, n(MY1)=11. **E)** Representative calcium transients under 0.5 Hz electrical stimulation (first five transients) and caffeine pulse (sixth transient) in cardiomyocytes isolated from MYBPC3-Q1061X-carrying EHTs (MY1) and their isogenic control cardiomyocytes (MY1-Ctrl). **F)** Kinetics of calcium transient upstroke and decay. n(MY1-Ctrl)=88, n(MY1)=115.**G)-H)** Amplitude (G) and decay (H) derived from electrically invoked calcium transients. n(MY1-Ctrl)=88, n(MY1)=115. **I)-J)** Amplitude (I) and decay (J) from caffeine invoked calcium transient n(MY1-Ctrl)=88, n(MY1)=115. **K)** Fractional calcium release assessed from the ratio of electrically and caffeine invoked **L)-O)** Representative traces (L), voltage-step protocol (M), current-voltage relationship (N), and voltage-dependent activation and inactivation (O) of L-type calcium current. In M and O n(MY1-Ctrl)=36, n(MY1)=27. Student’s *t*-test *P* value: *, *P* < 0.05.

**Supplementary Figure 4.**
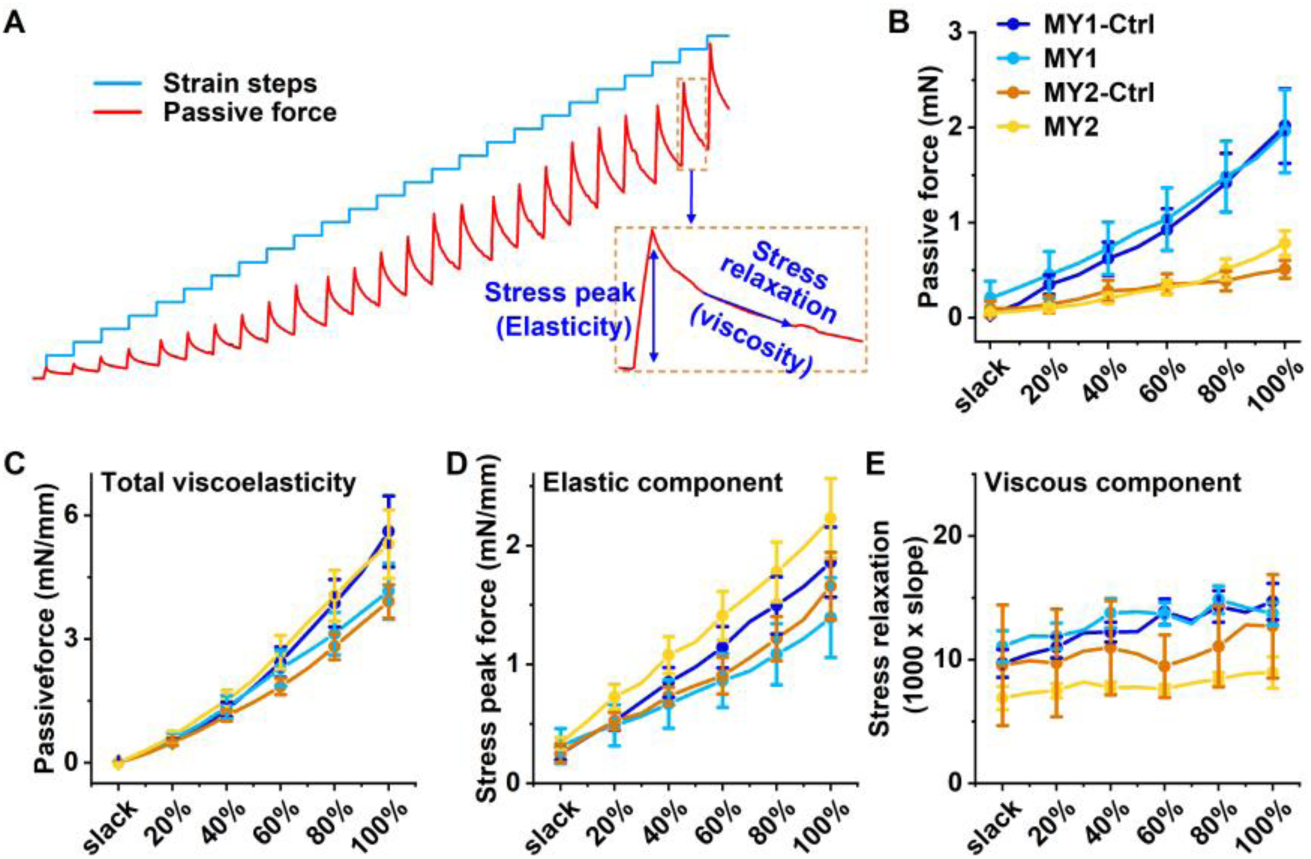
The passive and the viscoelastic properties of the EHTs. **A)** Representative passive force trace from isometric EHT force recording (red) illustrated with strain steps (blue) from the same recording. Inset highlights how different passive and viscoelastic parameters were assessed from the trace. **B)-E)** Passive force (B), total viscoelasticity (C), elasticity (D) and viscosity (E) assessed from the passive force traces. n(MY1-Ctrl)=32, n(MY1)=28, n(MY2-Ctrl)=20, n(MY2)=25.

**Supplementary Figure 5.**
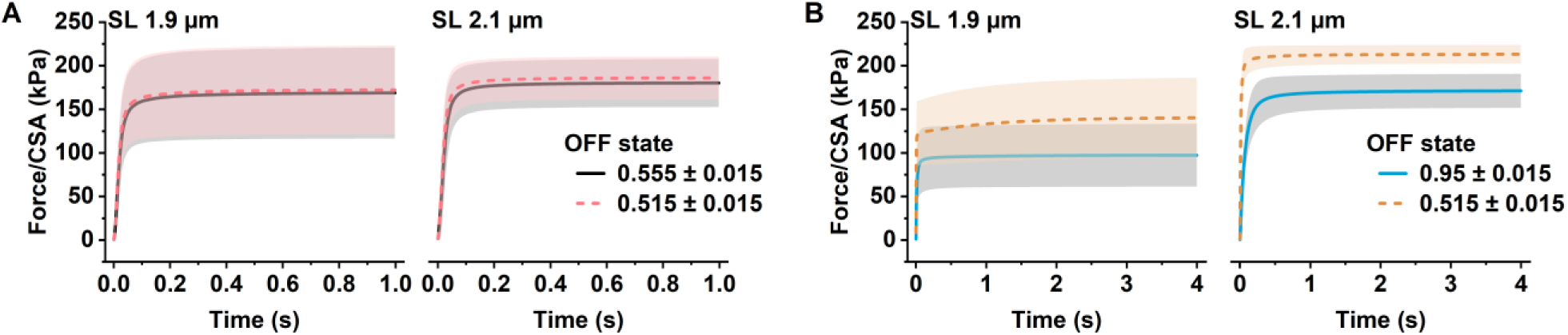
Simulated steady-state forces with cooperativity **A)** and without cooperativity **B)**.

**Supplementary Figure 6.**
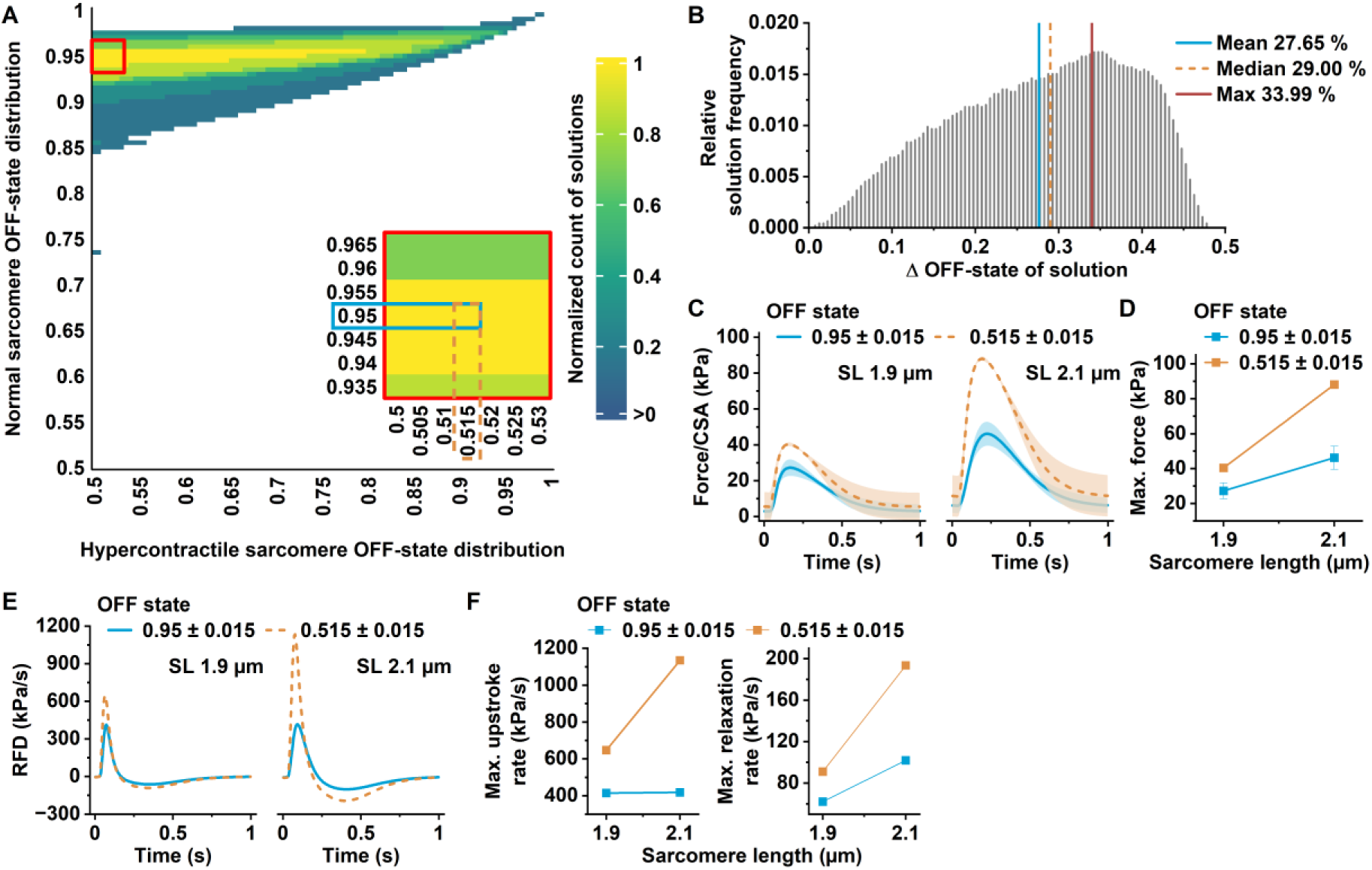
Simulations without cooperativity require a larger shift in ON/OFF ratio to reproduce hypercontractility, leading to slower relaxation and high resting tension. **A)** Heatmap of myosin OFF-state distributions in normal and hypercontractile sarcomeres. Color indicates the normalized number of model solutions for each distribution combination that reproduce the force changes observed in EHT experiments. The inset highlights the region of solutions used for the force plots in panel C. **B)** Distribution of myosin OFF-state differences across solutions. Note that the solutions are spread over a wider range. **C)** Isometric twitch forces at two sarcomere lengths. **D)** Maximum isometric forces at two sarcomere lengths, obtained from panel C. **E)** Rate of force development (RFD) during isometric twitch at different sarcomere lengths acquired by derivatization of isometric contraction traces in panel C. **F)** Isometric contraction maximum upstroke and relaxation rates at different sarcomere lengths determined from RFD plots in panel E.

**Supplementary Figure 7.**
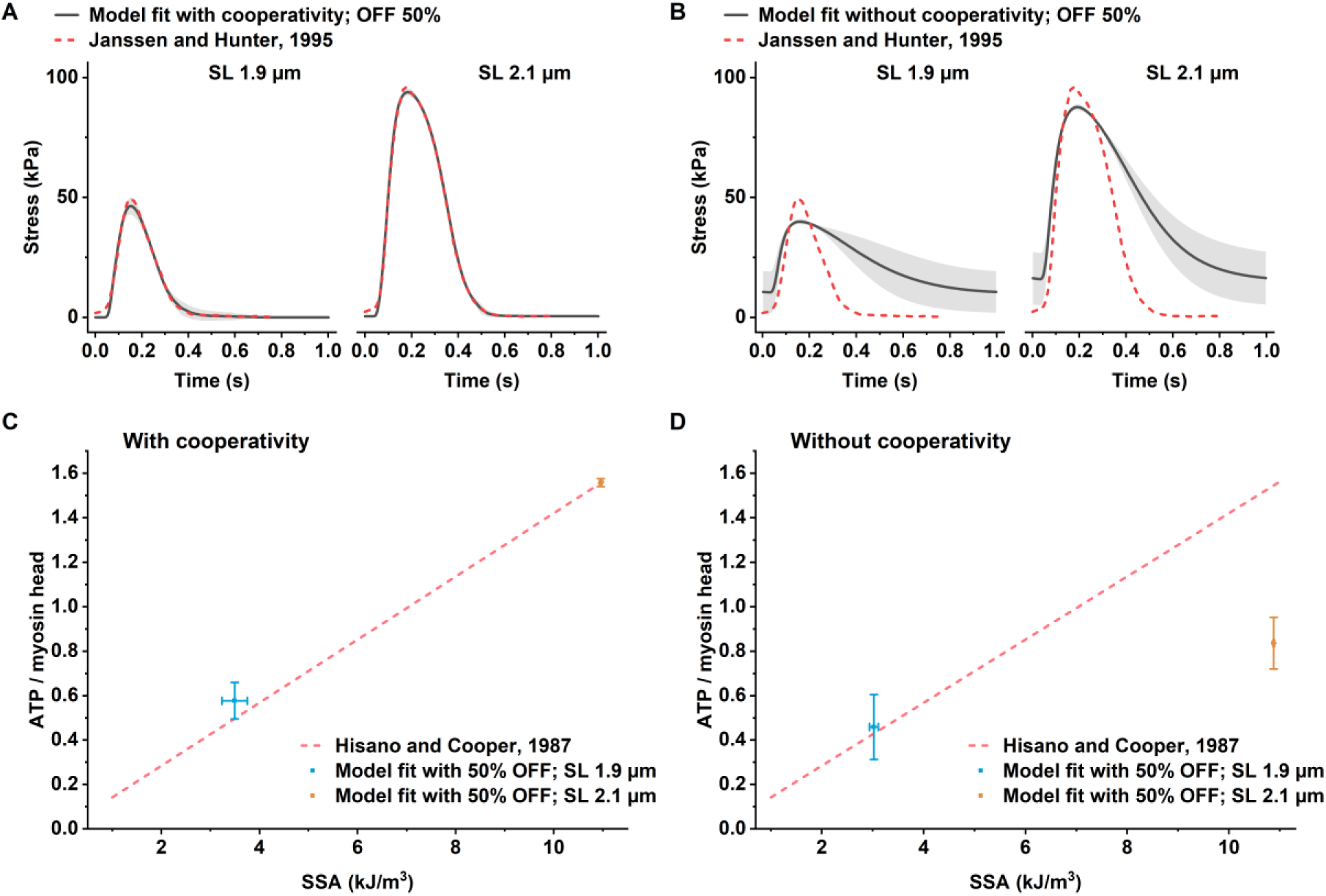
Model optimization results for selected parameter sets used in subsequent simulations with varying myosin OFF states. Fits to isometric stress data from rat trabeculae (Janssen and Hunter, 1995) with cooperativity **A)** and without cooperativity **B)**. Fits to the linear relationship between oxygen consumption and the stress–strain area (Hisano and Cooper, 1987) with cooperativity **C)** and without cooperativity **D)**. For simulations including cooperativity, a total of 256 optimizations were performed at each of the two sarcomere lengths using different combinations of free energy parameters. From these, 49 parameter sets were selected based on minimal residuals in additional simulations, with the constraint that energy parameters were identical for both sarcomere lengths. In the absence of cooperativity, 16 optimizations were performed, all of which were retained for further simulations. In this case, two tropomyosin free energy parameters were fixed to zero, reducing the number of parameter combinations and consequently, the total number of optimizations. Results are presented as mean ± SD.

**Supplementary Table 1.** Linear mixed effect model statistics for EHT Frank-Starling data.

| Linear mixed effect model: Dependant variable ~ Stretch * Patient * Variant + (1 \| Tissue) |  |  |  |  |  |  |  |  |  |  |  |  |  |
| --- | --- | --- | --- | --- | --- | --- | --- | --- | --- | --- | --- | --- | --- |
| Dependant variable | Effect of variant in each stretch level within patient |  |  |  |  |  |  | Test for the model's interaction contrasts within patient |  |  |  |  |  |
|  | Stretch | Patient | Estimate | SE | df | t.ratio | p.value | Patient | Effect | df1 | df2 | F.ratio | p.value |
| Twitch force |  |  |  |  |  |  |  |  |  |  |  |  |  |
|  | Slack | MY1 | -48.8 | 40.1 | 91.8 | -1.22 | 1.000 | MY1 | Stretch | 5 | 355 | 109 | <0.001 |
|  |  | MY2 | -60.8 | 44.4 | 91.8 | -1.37 | 1.000 |  | Variant | 1 | 71 | 11.7 | 0.001 |
|  | 20 % | MY1 | -112 | 40.1 | 91.8 | -2.79 | 0.607 |  | Stretch:Variant (interaction) | 5 | 355 | 8.58 | <0.001 |
|  |  | MY2 | -164 | 44.4 | 91.8 | -3.70 | 0.035 |  |  |  |  |  |  |
|  | 40 % | MY1 | -128 | 40.1 | 91.8 | -3.20 | 0.181 | MY2 | Stretch | 5 | 355 | 92.5 | <0.001 |
|  |  | MY2 | -227 | 44.4 | 91.8 | -5.12 | <0.001 |  | Variant | 1 | 71 | 24.6 | <0.001 |
|  | 60 % | MY1 | -146 | 40.1 | 91.8 | -3.64 | 0.044 |  | Stretch:Variant (interaction) | 5 | 355 | 22.9 | <0.001 |
|  |  | MY2 | -244 | 44.4 | 91.8 | -5.49 | 0.000 |  |  |  |  |  |  |
|  | 80 % | MY1 | -162 | 40.1 | 91.8 | -4.05 | 0.010 |  |  |  |  |  |  |
|  |  | MY2 | -260 | 44.4 | 91.8 | -5.85 | <0.001 |  |  |  |  |  |  |
|  | 100 % | MY1 | -174 | 40.1 | 91.8 | -4.34 | 0.004 |  |  |  |  |  |  |
|  |  | MY2 | -281 | 44.4 | 91.8 | -6.32 | <0.001 |  |  |  |  |  |  |
| Max. upstroke rate |  |  |  |  |  |  |  |  |  |  |  |  |  |
|  | Slack | MY1 | -0.29 | 0.224 | 85.7 | -1.30 | 1.000 | MY1 | Stretch | 5 | 344 | 108 | <0.001 |
|  |  | MY2 | -0.54 | 0.260 | 86.2 | -2.06 | 1.000 |  | Variant | 1 | 69 | 9.26 | 0.003 |
|  | 20 % | MY1 | -0.58 | 0.224 | 85.7 | -2.58 | 1.000 |  | Stretch:Variant (interaction) | 5 | 344 | 6.23 | <0.001 |
|  |  | MY2 | -1.16 | 0.260 | 85.7 | -4.45 | 0.002 |  |  |  |  |  |  |
|  | 40 % | MY1 | -0.66 | 0.224 | 85.7 | -2.93 | 0.412 | MY2 | Stretch | 5 | 344 | 92.4 | <0.001 |
|  |  | MY2 | -1.47 | 0.260 | 85.7 | -5.66 | <0.001 |  | Variant | 1 | 69 | 30.0 | <0.001 |
|  | 60 % | MY1 | -0.72 | 0.224 | 85.7 | -3.20 | 0.184 |  | Stretch:Variant (interaction) | 5 | 344 | 23.2 | <0.001 |
|  |  | MY2 | -1.52 | 0.260 | 85.7 | -5.87 | <0.001 |  |  |  |  |  |  |
|  | 80 % | MY1 | -0.76 | 0.224 | 85.7 | -3.41 | 0.094 |  |  |  |  |  |  |
|  |  | MY2 | -1.64 | 0.260 | 85.7 | -6.32 | <0.001 |  |  |  |  |  |  |
|  | 100 % | MY1 | -0.86 | 0.224 | 85.7 | -3.86 | 0.021 |  |  |  |  |  |  |
|  |  | MY2 | -1.75 | 0.260 | 85.7 | -6.75 | <0.001 |  |  |  |  |  |  |
| Max. relaxation rate |  |  |  |  |  |  |  |  |  |  |  |  |  |
|  | Slack | MY1 | -0.18 | 0.127 | 85.3 | -1.46 | 1.000 | MY1 | Stretch | 5 | 337 | 108 | <0.001 |
|  |  | MY2 | -0.26 | 0.152 | 87.0 | -1.72 | 1.000 |  | Variant | 5 | 337 | 69.1 | <0.001 |
|  | 20 % | MY1 | -0.34 | 0.127 | 85.3 | -2.65 | 0.923 |  | Stretch:Variant (interaction) | 1 | 67 | 11.5 | 0.001 |
|  |  | MY2 | -0.41 | 0.151 | 85.3 | -2.72 | 0.752 |  |  |  |  |  |  |
|  | 40 % | MY1 | -0.40 | 0.127 | 85.3 | -3.13 | 0.231 | MY2 | Stretch | 1 | 68 | 14.3 | <0.001 |
|  |  | MY2 | -0.56 | 0.151 | 85.3 | -3.74 | 0.032 |  | Variant | 5 | 337 | 8.73 | <0.001 |
|  | 60 % | MY1 | -0.44 | 0.127 | 85.3 | -3.48 | 0.077 |  | Stretch:Variant (interaction) | 5 | 337 | 9.68 | <0.001 |
|  |  | MY2 | -0.61 | 0.151 | 85.3 | -4.03 | 0.012 |  |  |  |  |  |  |
|  | 80 % | MY1 | -0.51 | 0.127 | 85.3 | -4.04 | 0.011 |  |  |  |  |  |  |
|  |  | MY2 | -0.68 | 0.151 | 85.3 | -4.49 | 0.002 |  |  |  |  |  |  |
|  | 100 % | MY1 | -0.56 | 0.127 | 85.3 | -4.46 | 0.002 |  |  |  |  |  |  |
|  |  | MY2 | -0.72 | 0.151 | 85.3 | -4.74 | <0.001 |  |  |  |  |  |  |

**Supplementary Table 2.** Linear mixed effect model statistics for EHT frequency protocol data.

| Linear mixed effect model: Dependant variable ~ Frequency * Patient * Variant + (1 \| Tissue) |  |  |  |  |  |  |  |  |  |  |  |  |  |
| --- | --- | --- | --- | --- | --- | --- | --- | --- | --- | --- | --- | --- | --- |
| Dependant variable | Effect of variant in each pacing frequency within patient |  |  |  |  |  |  | Test for the model's interaction contrasts within patient |  |  |  |  |  |
|  | Freq. | Patient | Estimate | SE | df | t.ratio | p.value | Patient | Effect | df1 | df2 | F.ratio | p.value |
| **Baseline force** |  |  |  |  |  |  |  |  |  |  |  |  |  |
|  | 0.5 Hz | MY1 | 10.1 | 15.0 | 181 | 0.68 | 1 | MY1 | Frequency | 3 | 157 | 33.2 | <0.001 |
|  |  | MY2 | 8.76 | 18.0 | 185 | 0.49 | 1 |  | Variant | 1 | 61 | 5.89 | 0.018 |
|  | 1 Hz | MY1 | 11.2 | 14.6 | 176 | 0.77 | 1 |  | Freq.:Variant (interaction) | 3 | 157 | 10.2 | <0.001 |
|  |  | MY2 | 5.80 | 19.1 | 196 | 0.30 | 1 |  |  |  |  |  |  |
|  | 2 Hz | MY1 | -42.5 | 15.2 | 183 | -2.81 | 0.266 | MY2 | Frequency | 3 | 158 | 41.3 | <0.001 |
|  |  | MY2 | -42.9 | 19.5 | 199 | -2.20 | 1 |  | Variant | 1 | 63 | 8.64 | 0.005 |
|  | 3 Hz | MY1 | -79.5 | 17.1 | 202 | -4.66 | <0.001 |  | Freq.:Variant (interaction) | 3 | 158 | 15.1 | <0.001 |
|  |  | MY2 | -115 | 17.2 | 176 | -6.72 | <0.001 |  |  |  |  |  |  |
| **Twitch force** |  |  |  |  |  |  |  |  |  |  |  |  |  |
|  | 0.5 Hz | MY1 | -137 | 36.7 | 69.2 | -3.74 | 0.018 | MY1 | Frequency | 3 | 152 | 43.9 | <0.001 |
|  |  | MY2 | -220 | 43.1 | 69.8 | -5.09 | <0.001 |  | Variant | 1 | 56 | 8.17 | 0.006 |
|  | 1 Hz | MY1 | -139 | 36.7 | 69.2 | -3.80 | 0.015 |  | Freq.:Variant (interaction) | 3 | 152 | 10.9 | <0.001 |
|  |  | MY2 | -220 | 43.8 | 73.8 | -5.01 | <0.001 |  |  |  |  |  |  |
|  | 2 Hz | MY1 | -90.7 | 36.7 | 69.2 | -2.47 | 0.760 | MY2 | Frequency | 3 | 152 | 25.4 | <0.001 |
|  |  | MY2 | -159 | 43.3 | 71.0 | -3.66 | 0.023 |  | Variant | 1 | 56 | 17.6 | <0.001 |
|  | 3 Hz | MY1 | -30.0 | 37.9 | 78.0 | -0.79 | 1 |  | Freq.:Variant (interaction) | 3 | 152 | 12.9 | <0.001 |
|  |  | MY2 | -86.9 | 43.7 | 73.3 | -1.99 | 1 |  |  |  |  |  |  |
| **Max. upstroke rate** |  |  |  |  |  |  |  |  |  |  |  |  |  |
|  | 0.5 Hz | MY1 | -0.83 | 0.218 | 58.8 | -3.78 | 0.017 | MY1 | Frequency | 3 | 144 | 46.1 | <0.001 |
|  |  | MY2 | -1.35 | 0.250 | 59.3 | -5.40 | <0.001 |  | Variant | 1 | 54 | 9.96 | 0.003 |
|  | 1 Hz | MY1 | -0.85 | 0.218 | 58.8 | -3.91 | 0.012 |  | Freq.:Variant (interaction) | 3 | 144 | 14.7 | <0.001 |
|  |  | MY2 | -1.35 | 0.251 | 60.0 | -5.36 | <0.001 |  |  |  |  |  |  |
|  | 2 Hz | MY1 | -0.69 | 0.218 | 59.1 | -3.15 | 0.123 | MY2 | Frequency | 3 | 144 | 29.4 | <0.001 |
|  |  | MY2 | -1.15 | 0.251 | 59.7 | -4.59 | 0.001 |  | Variant | 1 | 54 | 23.1 | <0.001 |
|  | 3 Hz | MY1 | -0.33 | 0.222 | 63.0 | -1.49 | 1 |  | Freq.:Variant (interaction) | 3 | 144 | 13.4 | <0.001 |
|  |  | MY2 | -0.85 | 0.251 | 59.7 | -3.39 | 0.061 |  |  |  |  |  |  |
| **Max. relaxation rate** |  |  |  |  |  |  |  |  |  |  |  |  |  |
|  | 0.5 Hz | MY1 | -0.35 | 0.125 | 60.4 | -2.77 | 0.354 | MY1 | Stretch | 3 | 118 | 20.4 | <0.001 |
|  |  | MY2 | -0.48 | 0.125 | 62.8 | -3.80 | 0.016 |  | Variant | 1 | 48 | 11.8 | 0.001 |
|  | 1 Hz | MY1 | -0.33 | 0.125 | 60.4 | -2.59 | 0.570 |  | Stretch:Variant (interaction) | 3 | 118 | 2.26 | 0.085 |
|  |  | MY2 | -0.48 | 0.128 | 68.7 | -3.77 | 0.017 |  |  |  |  |  |  |
|  | 2 Hz | MY1 | -0.50 | 0.127 | 62.4 | -3.96 | 0.009 | MY2 | Stretch | 3 | 118 | 27.9 | <0.001 |
|  |  | MY2 | -0.77 | 0.124 | 60.4 | -6.24 | <0.001 |  | Variant | 1 | 47 | 31.0 | <0.001 |
|  | 3 Hz | MY1 | -0.45 | 0.140 | 88.4 | -3.22 | 0.086 |  | Stretch:Variant (interaction) | 3 | 118 | 12.5 | <0.001 |
|  |  | MY2 | -0.85 | 0.124 | 60.4 | -6.88 | <0.001 |  |  |  |  |  |  |

## Notes

### Competing Interest Statement

The authors have declared no competing interest.

